# Methylated guanosine and uridine modifications in *S. cerevisiae* mRNAs modulate translation elongation

**DOI:** 10.1101/2022.06.13.495843

**Authors:** Joshua D. Jones, Monika K. Franco, Tyler J. Smith, Laura R. Snyder, Anna G. Anders, Brandon T. Ruotolo, Robert T. Kennedy, Kristin S. Koutmou

## Abstract

Chemical modifications to protein encoding messenger RNA (mRNA) can modulate their localization, translation and stability within cells. Over 15 different types of mRNA modifications have been identified by sequencing and liquid chromatography coupled to tandem mass spectrometry (LC-MS/MS) technologies. While LC-MS/MS is arguably the most essential tool available for studying analogous protein post-translational modifications, the high-throughput discovery and quantitative characterization of mRNA modifications by LC-MS/MS has been hampered by the difficulty of obtaining sufficient quantities of pure mRNA and limited sensitivities for modified nucleosides. To overcome these challenges, we improved the mRNA purification and LC-MS/MS pipelines to identify new *S. cerevisiae* mRNA modifications and quantify 50 ribonucleosides in a single analysis. The methodologies we developed result in no detectable non-coding RNA modifications signals in our purified mRNA samples and provide the lowest limit of detection reported for ribonucleoside modification LC-MS/MS analyses. These advancements enabled the detection and quantification of 13 *S. cerevisiae* mRNA ribonucleoside modifications and revealed four new *S. cerevisiae* mRNA modifications at low to moderate levels (1-methyguanosine, N2-methylguanosine, N2, N2-dimethylguanosine, and 5-methyluridine). We identified four enzymes that incorporate these modifications into *S. cerevisiae* mRNAs (Trm10, Trm11, Trm1, and Trm2), though our results suggest that guanosine and uridine nucleobases are also non-enzymatically methylated at low levels. Regardless of whether they are incorporated in a programmed manner or as the result of RNA damage, we reasoned that the ribosome will encounter the modifications that we detect in cells and used a reconstituted translation system to discern the consequences of modifications on translation elongation. Our findings demonstrate that the introduction of 1-methyguanosine, N2-methylguanosine and 5-methyluridine into mRNA codons impedes amino acid addition in a position dependent manner. This work expands the repertoire of nucleoside modifications that the ribosome must decode in *S. cerevisiae*. Additionally, it highlights the challenge of predicting the effect of discrete modified mRNA sites on translation *de novo* because individual modifications influence translation differently depending on mRNA sequence context.

## INTRODUCTION

Post-transcriptional modifications to RNA molecules can change their structure, localization, stability, and function^1,2^. To date, over 150 different nucleoside chemical modifications have been identified within non-coding RNAs (ncRNA), and many are important, or even essential, for a myriad of cellular processes^1,3^. The significance of RNA modifications to cellular health is underscored by decades of observations implicating the mis-regulation of ncRNA modifying enzymes in cancer and other diseases^4–9^. Recent advances in next generation RNA sequencing (RNA-seq)^10–19^ and liquid chromatography coupled to tandem mass spectrometry (LC-MS/MS) technologies^20–24^ enabled the detection of chemical modifications in protein encoding messenger RNAs (mRNA). Over 15 mRNA modifications have been reported, including N6-methyladensoine (m^6^A), inosine (I), N7-methylguanosine (m^7^G), and pseudouridine (Ψ)^1,12,13,22,25–28^. There are >10-fold more types of modifications reported in ncRNA than in mRNA, raising the possibility that the diversity of mRNA modifications has not yet been revealed.

While the biological significance of ncRNA modifications has been extensively studied, the consequences of mRNA modifications on gene expression are just beginning to be explored. Modified nucleosides resulting from RNA damage (e.g. oxidation, alkylation, or UV) commonly perturb protein synthesis and can trigger RNA degradation pathways^29,30^. Despite typically being present at lower levels than their enzymatically incorporated counterparts^31^, there is evidence that oxidized mRNAs can accumulate in neurodegenerative diseases^31–33^. The most abundant and well-studied modification added by enzymes into mRNA coding regions, m^6^A, has been implicated as a key modulator of multiple facets of the mRNA lifecycle including nuclear export^34–36^, mRNA stability^37–39^, and translational efficiency^19,38,40–43^. Given these potential contributions to mRNA function, it is unsurprising that the mis-regulation of m^6^A is linked to a host of diseases such as endometrial cancer^44^ and type 2 diabetes^45^. While initial studies of m^6^A provide an example of the biological impact mRNA modifications can have, most other mRNA modifications have been minimally investigated. The development of additional sensitive and quantitative techniques to comprehensively evaluate the mRNA modification landscape will be essential to direct future investigations that characterize the molecular level consequence of emerging mRNA modifications.

LC-MS/MS has been a powerful approach to characterize chemical modifications of all three major classes of bio-molecules central to protein synthesis (DNA, RNA, and protein). In particular, the sensitivity and specificity of LC-MS/MS methodologies have enabled the identification and extensive characterization of post-translational protein modifications^46^. While post-transcriptional modifications of ncRNA have been studied for decades using 2D thin layer chromatography^47^ and LC coupled to ultraviolet detection^48,49^, recent developments in LC-MS/MS analyses provided some of the first insight into RNA modification abundance and dynamics under cellular stress^50–56^. Such methods can broadly detect and provide absolute quantification of modifications in any purified RNA sample^25^. These features have made LC-MS/MS an attractive technology to adopt for mRNA modification discovery. Currently, published methods can assay up to 40 ribonucleosides in a single analysis and use calibration curves from standards to enable quantification with high accuracy and selectivity^20^. However, despite these advantages and the proven utility of LC-MS/MS methodologies for investigating ncRNA modifications, LC-MS/MS has yet to be widely used to study mRNA modifications unlike the comprehensive characterization of post-translational protein modifications by LC-MS/MS technologies over the past few decades.

Here, we identify two factors that have impeded application of LC-MS/MS to mRNA modification analysis: the quantity of mRNA required for current LC-MS/MS sensitivities, and the difficulty to obtain highly pure mRNA. We integrated an improved chromatographic approach with an enhanced mRNA purification and validation process to overcome these limitations and develop a robust workflow for mRNA modification characterization. Our method is capable of quantifying 50 ribonucleoside variants in a single analysis. Analysis of purified *S. cerevisiae* mRNA samples reveals that 1-methylguanosine (m^1^G), N2-methylguanosine (m^2^G), N2, N2-dimethylguanosine (m^2^_2_G), and 5-methyluridine (m^5^U) are likely incorporated into mRNAs both enzymatically (Trm10, Trm11, Trm1, and Trm2) and non-enzymatically. We also use a fully purified *in vitro* translation system to demonstrate that the inclusion of these methylated nucleosides into mRNA codons can slow amino acid addition by the ribosome. Together, our findings advance available chromatography and mRNA purification and validation methods to enhance the high-confidence and high-throughput detection of modified nucleosides by LC-MS/MS and support a growing body of evidence that the inclusion of mRNA modifications commonly alters the peptide elongation during protein synthesis.

## RESULTS AND DISCUSSION

### Development of highly sensitive LC-MS/MS method for simultaneously quantifying 50 ribonucleosides

RNA-seq based technologies capable of identifying the location of RNA modifications have revealed that modified nucleosides can be found in thousands of mRNAs^57^. These powerful methodologies have enabled the widespread study of mRNA modifications, but are computationally laborious, not generally quantitative, and typically detect a single modification at a time. In contrast, LC-MS/MS analyses rapidly and quantitatively identify the presence of RNA modifications but cannot report on where they exist throughout the transcriptome^25^. Therefore, the integration of orthogonal LC-MS/MS and RNA-seq based methodologies is required to develop robust platforms for detecting mRNA modifications^57–66^. However, the application of LC-MS/MS for nucleoside discovery has been limited by lingering questions regarding mRNA purity, as many reports do not present the comprehensive quality controls necessary for confident mRNA modification analysis. Indeed, a few reported mRNA modifications have not yet been mapped to discrete mRNAs in the transcriptome by RNA-seq based methodologies (e.g., m^1^G), likely due to their low abundance and/or possible non-specific incorporation. While there is evidence that the insertion of some mRNA modifications are programmed, suggesting a biological function, other modifications are likely added in a less specific manner (e.g., RNA damage, off target modification by ncRNA enzymes). Modifications incorporated at lower levels are unlikely to be detected by sequencing-based methods, but can have consequences for cellular health, as illustrated by links between RNA-damage and disease. Therefore, regardless of why a modification is present, it is still essential for us to fully elucidate the mRNA modification landscape and interrogate how these modifications affect cellular function.

Ribonucleosides are most commonly separated using reversed phase chromatography and quantified using multiple reaction monitoring (MRM) on a triple quadrupole mass spectrometer^20,50,52,67,68^. These methods have reported limits of detection (LODs) down to ∼60 attomole for select ribonucleosides using standard mixtures with canonical and modified nucleosides at equal concentrations^50^. However, the abundance of unmodified and modified nucleosides in RNAs are not equivalent in cells, with canonical bases existing in 20-to 10,000-fold higher concentrations than RNA modifications **(Figure 1A)**. In currently available chromatography methods, modified nucleosides (e.g., m^5^U, m^1^G, m^1^Ψ, and s^2^U) commonly coelute with canonical nucleosides, reducing the detectability of some modified bases^50,52,53^. Coelution limits the utility of available LC-MS/MS methods because it results in ion suppression of modified nucleoside signals, with abundant canonical nucleosides outcompeting modified nucleosides for electrospray droplet surface charge. Additionally, this phenomenon makes calibration curves non-linear and worsens the quantifiability of modifications at concentrations necessary for mRNA modification analyses. Recent efforts have been made to derivatize ribonucleosides prior to LC-MS/MS analysis to increase sensitivity and retention on reversed-phase chromatography^21,69–71^. The analogous benzoyl chloride derivatization of neurochemicals has previously been an important separation strategy for many neurochemical monitoring applications^72,73^. However, labeling strategies are unlikely to prove as useful for investigating mRNA modifications because derivatizing agents are typically nucleobase specific, limiting the ability of LC/MS-MS assays to be multiplexed^21,69,70^. Furthermore, labeling increases the amount of mRNA sample required due to additional sample preparation steps following derivatization. This is an important consideration given that mRNAs represent only ∼1-2% of the total RNAs in a cell, and it is already challenging to purify sufficient quantities of mRNA for LC-MS/MS analysis.

**Figure 1:**
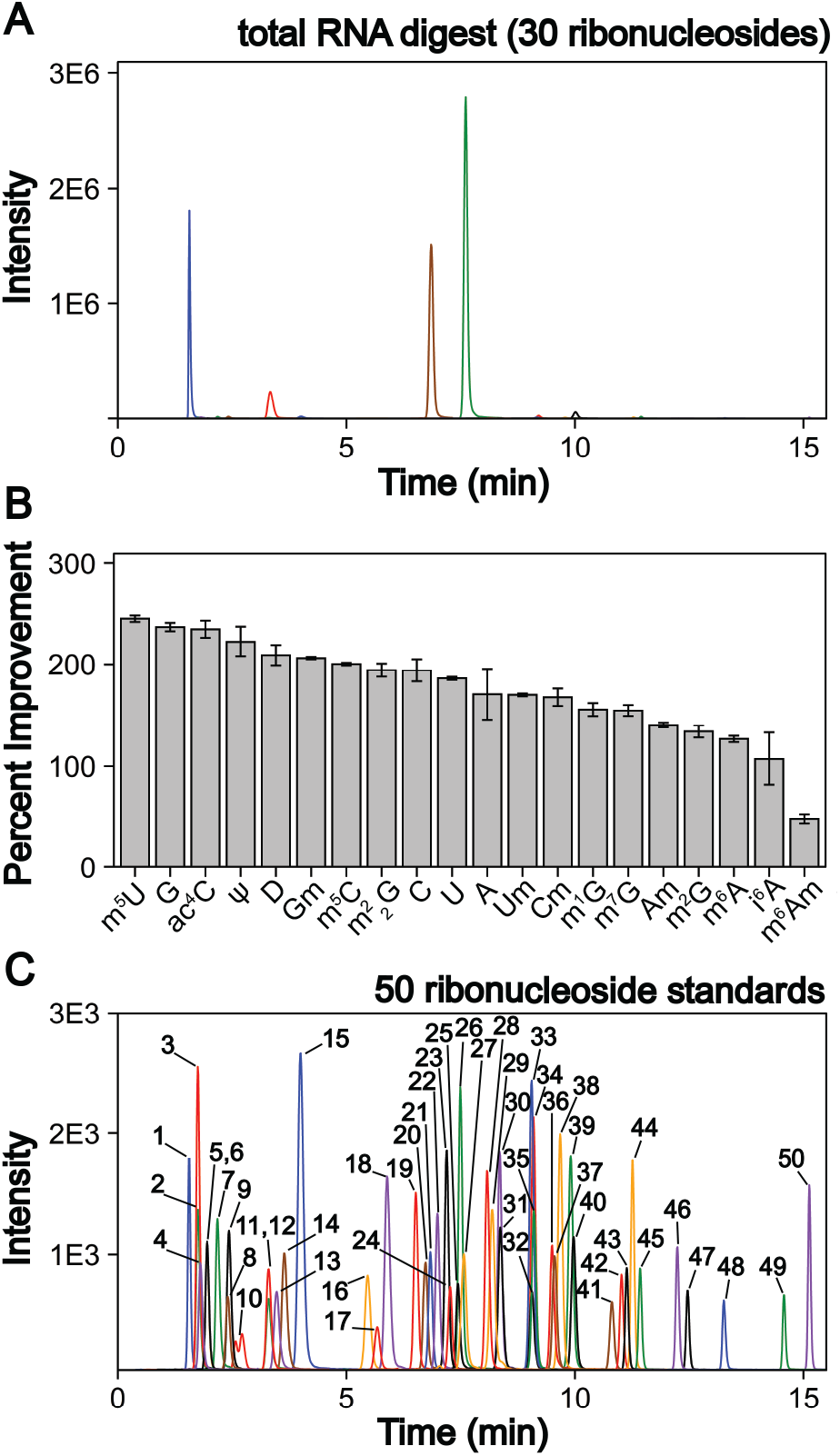
LC-MS/MS method development to quantify 50 ribonucleosides in a single analysis. **A)** Extracted ion chromatogram for the 30 ribonucleosides (4 canonical bases and 26 naturally occurring modifications) detected in a *S. cerevisiae* total RNA digestion displaying that the canonical bases exist at much larger levels than the ribonucleoside modifications. **B)** LC-MS/MS signal percent improvement using 1 mm chromatography at 100 μL/min compared to 2 mm chromatography at 400 μL/min. **C)** Extracted ion chromatogram for 50 ribonucleoside standards (4 canonical bases, 45 naturally occurring modifications, and 1 non-natural modifications). The concentrations of each ribonucleoside standards within the standard mix and their corresponding peak numbers are displayed in **Supplemental Table S2**. For the chromatograms, each color peak represents a separate ribonucleoside in the method, and the colors are coordinated between panel A and C.

We addressed these limitations by first improving upon existing chromatography techniques. Current methods typically utilize 2 mm internal diameter (I.D.) columns that require higher flow rates (300 to 400 μL/min), which worsens ionization efficiencies than smaller I.D. chromatography with lower flow rates. We utilized a 1 mm I.D. column with flow rates at 100 μL/min to lessen these effects. In principle, even smaller bore columns (i.e., “nano-LC”), which are commonly used in in proteomics^74^, could be used. Indeed, some studies have shown their effectiveness for nucleosides^75,76^; however, smaller bore columns can suffer from robustness issues in some conditions. Also, low binding capacity of more polar nucleosides results in poor peak shapes in nano-LC because of relatively large injection volumes. Another limitation has been the stationary phases used, where porous graphitic carbon columns yield poor chromatographic performance for some ribonucleosides (e.g., methylated guanosine modifications) and many C18 phases have low binding capacity for some ribonucleosides (e.g., cytidine and pseudouridine) making them difficult to retain. We used a polar endcapped C18 column to provide more retention and good performance for all nucleosides. We also used mobile phase buffers which have previously been shown to provide high ESI-MS sensitivity for modified ribonucleosides^50^. These alterations combined increased the sensitivity of the assay by 50 to 250% for all nucleosides tested compared to standard 2 mm I.D. chromatography at 400 μL/min **(Figure 1B)** while maintaining adequate ribonucleoside binding capacity for early eluting ribonucleosides. We also altered the chromatographic conditions including increased temperature (35°C vs 25°C) and modified mobile phase gradients to prevent coelution of the highly abundant canonical nucleosides with the modified nucleosides. Notably, in contrast to most available methods, m^5^U, m^1^G, m^1^Ψ do not coelute with unmodified nucleosides in our method **(Figure 1C)**. This improved separation greatly reduced ionization suppression of these nucleosides. Together, these advancements led to a wider linear dynamic range than previous reports with over four orders of magnitude for most modifications and LODs down to 3 amol (0.6 pM) using a single internal standard and no derivatization steps. Our method represents at least a 10-fold improvement over previous ultrahigh-performance LC (UHPLC) and nano-LC analyses for most modifications analyzed **(Supplemental Table S1 and Supplemental Figures S1 through S4)**. Therefore, the method described here provides a linear dynamic range and LODs capable of analyzing both highly modified ncRNA in addition to the less modified mRNA without large sample requirements. To perform an in-depth RNA modification analysis, approximately 50 to 200 ng of total RNA or mRNA is required per replicate which is achievable using standard eukaryotic and bacterial cell culture techniques. Overall, this assay can quantify the 4 canonical nucleosides, 45 naturally occurring modified nucleosides, and 1 non-natural modified nucleoside (internal control) **(Figure 1C, Supplemental Table S2)**. This work ameliorates current quantitative ribonucleoside LC-MS/MS methodologies by improving chromatographic conditions and characterizing quantifiability at nucleoside concentrations representative of typical RNA digest samples to enable higher confidence total RNA and mRNA modification analyses.

### Three-stage mRNA purification and validation pipeline provides highly pure *S. cerevisiae* mRNA

Total RNA is mainly comprised of the highly modified transfer RNA (tRNA) and ribosomal RNA (rRNA) with a small percentage of mRNA. Unlike RNA-seq, LC-MS/MS assays are unable to distinguish between modifications arising from ncRNA or mRNA. In total RNA digestions, mRNA modifications typically exist at least 100X lower concentrations than in the corresponding total RNA samples^20^. Thus, even low-level contamination of tRNA and rRNA in purified mRNA samples can lead to inaccurate quantifications as well as false mRNA modifications discoveries. Most of the published mRNA purification pipelines use a combination of poly(A) enrichment and rRNA depletion steps to obtain mRNA^10,12,20,22,24,77,78^. However, previously this was found to be insufficient for removing all signal from contaminating ncRNA modifications during LC-MS/MS analyses, especially from contaminating tRNA^20,79^. The inability to obtain convincingly pure mRNA samples has long limited the utility of LC-MS/MS for studying these molecules. Recently, small RNA depletion steps have begun to be incorporated into mRNA purification pipelines to remove residual tRNA contamination^80^; however, the highest efficiency purifications typically require expensive instrumentation and materials (liquid chromatograph and size exclusion column)^23^ or expertise in RNA gel purification^21^. Despite these improvements, most reports do not provide adequate mRNA purity quality control to confirm removal of ncRNA for confident mRNA modification analyses. In order to apply our LC-MS/MS assay to studying mRNAs, we developed and implemented a three-stage purification pipeline comprised of a small RNA depletion step, two consecutive poly(A) enrichment steps, and ribosomal RNA depletion to selectively deplete the small ncRNA (e.g., tRNA and 5S rRNA) in addition to the 18S rRNA and 28S rRNA using fully commercial kits **(Figure 2)**. Additionally, we performed extensive quality control on our mRNA samples prior to LC-MS/MS analysis – assessing the purity of our mRNA following the three-stage purification pipeline using chip electrophoresis (Bioanalyzer), RNA-seq, qRT-PCR, and LC-MS/MS. The highly purified mRNA contained no detectable tRNA and rRNA peaks based on our Bioanalyzer electropherograms **(Figure 3A)**. The Bioanalyzer RNA 6000 pico assay provides an LOD of 25 pg/uL for a single RNA^81^; thus, the maximum theoretical tRNA or rRNA contamination would be 0.8% if it was just below our detection limit (3000 pg/uL sample analyzed). Similarly, RNA-seq indicated the mRNA is enriched from 4.1% in our total RNA to 99.8% in our purified mRNA samples **(Figure 3B, Supplemental Table S3)**. Additionally, we observed a >3000-fold depletion of 25S and 18S rRNAs and an >9-fold enrichment of actin mRNA based on qRT-PCR **(Supplemental Figure S5)**. Despite recent improvements in RNA-seq technologies and reverse transcriptases, the ability to accurately measure tRNA abundance by RNA-seq remains a struggle due to RNA modifications in these highly structured RNAs. While similar purities by RNA-seq have been achieved without a small RNA depletion step^20,78^, we previously found that this protocol was insufficient at removing all contaminating ncRNA signals by LC-MS/MS^20^ since RNA-seq does not accurately report on tRNA contamination^82^. While the RNA-seq could be improved by utilizing a more tRNA compatible reverse transcriptase, quality control analyses in addition to RNA-seq (such as LC-MS/MS) are necessary to judge tRNA contamination in purified mRNA.

**Figure 2:**
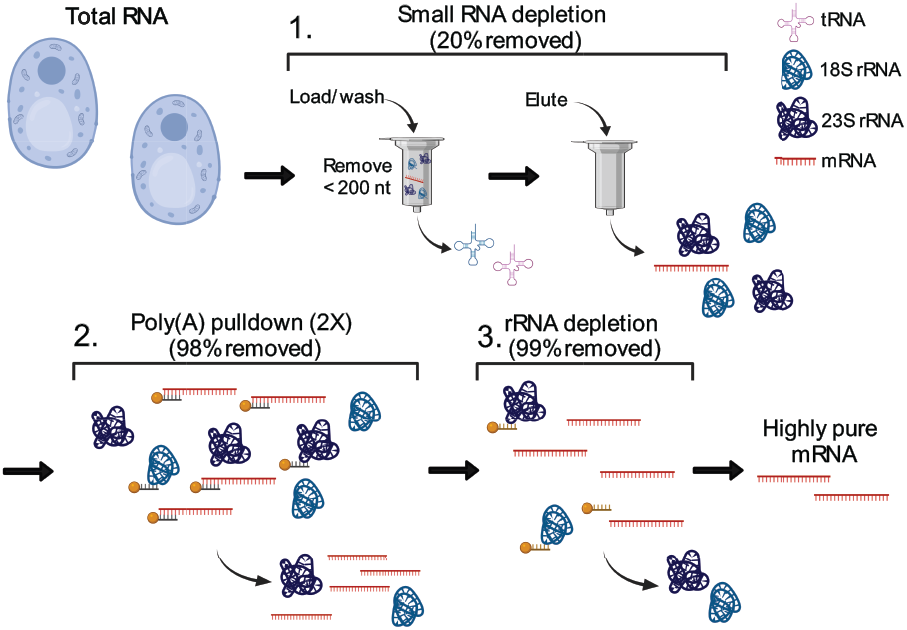
Three-stage mRNA purification pipeline. Total RNA from *S. cerevisiae* is purified to mRNA using a three-stage purification pipeline: 1. Small RNA (e.g., tRNA and 5S rRNA) is depleted; 2. mRNA is enriched from the small RNA depleted fraction through two consecutive poly(A) enrichment steps; 3. Remaining rRNA is depleted to result in highly purified mRNA. The displayed percent removed is the additive percent of total RNA removed throughout the three-stage purification pipeline.

**Figure 3:**
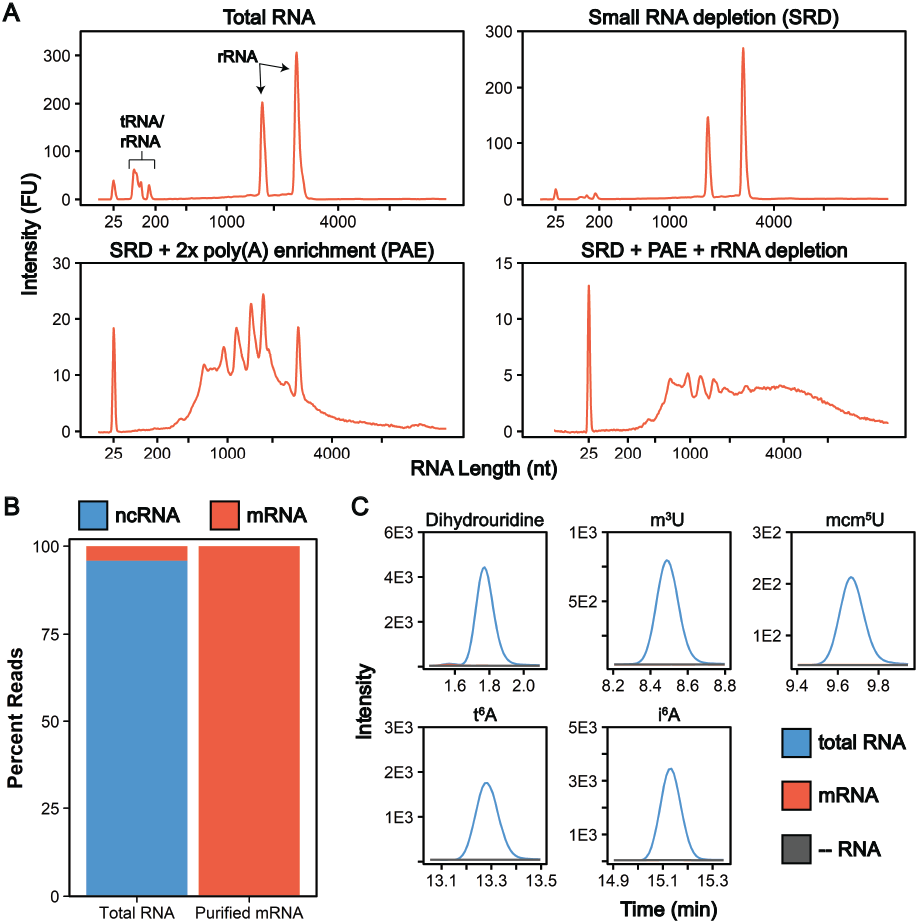
mRNA purity following three-stage purification pipeline. **A)** Bioanalyzer electropherograms displaying the RNA distribution following each stage of our purification pipeline. **B)** Average percentage of reads mapping to ncRNA (rRNA, tRNA, snRNA, etc.) and mRNA determined by RNA-seq of two biological replicate total RNA and purified mRNA samples. **C)** Representative overlaid extraction ion chromatograms for five RNA modifications that exist solely in ncRNA. These five modifications, in addition to eight additional ncRNA modifications, were detected in our total RNA samples (blue) while not detected in our mRNA samples (red) above our control digestions without RNA added (grey).

Since our highly multiplexed LC-MS/MS methodology is capable of quantifying known ncRNA and mRNA modifications in a single analysis, we can use this assay to further confirm the purity of our mRNA from the three-stage purification pipeline **(Figure 2)**. In these assays, total RNA and purified mRNA are degraded to ribonucleosides using a two-stage enzymatic digestion with Nuclease P1 and bacterial alkaline phosphatase **(Figure 4A)**. The resulting modified ribonucleosides are quantified and their concentrations are normalized to their corresponding canonical nucleosides (e.g., m^6^A/A) to account for variations in RNA quantities digested. In our total RNA samples, we detected 26 out of 30 known *S. cerevisiae* ribonucleoside modifications that we assayed for, where f^5^C, s^2^U, m^2,7^G, and m_3_G were not detected **(Figure 4B, Supplemental Table S4)**. This was expected because these modifications likely exist at levels below our LOD in our total RNA samples as they either arise from oxidative damage of m^5^C (f^5^C)^83,84^, are present at very low levels on *S. cerevisiae* tRNA (s^2^U)^85–87^, or are only found in low abundance snRNA and snoRNA (m^2,7^G and m_3_G)^88–90^. Additionally, we do not detect the 16 ribonucleoside modifications in our assay that have never been reported in *S. cerevisiae* (1 non-natural and 15 natural) **(Figure 4B, Supplemental Table S4)**. Our purified mRNA samples contained markedly fewer modifications than total RNA, as expected. In addition to the 16 non-*S. cerevisiae* modifications, we do not detect 13 *S. cerevisiae* non-coding RNA modifications that were present in our total RNA samples **(Figure 3C and Supplemental Table S4)**. All modifications not detected in the purified mRNA are reported to be exclusively located in *S. cerevisiae* tRNAs or rRNAs (e.g., i^6^A, m^3^C)^3^, result from oxidative damage (f^5^C)^64^, or were only previously detected in *S. cerevisiae* mRNAs purified from cells in grown under H_2_O_2_ stress (ac^4^C)^20^. The highly abundant dihydrouridine (DHU) modification provides a key example of such a common ncRNA modification that is not detected in our purified samples. DHU is located at multiple sites on every *S. cerevisiae* tRNA and is present at high levels (1.9 DHU/U%) in our total RNA samples **(Supplemental Table S5 and S6)**. However, we do not detect DHU above our LOD in our purified mRNA **(Figure 3C)**. Using this data, we estimated the maximum tRNA contamination in our purified mRNA to be 0.002% since DHU is not present in *S. cerevisiae* rRNAs **(Supplemental Calculation S1)**. The inability of our assay to detect highly abundant ncRNA modifications such as DHU provides orthogonal evidence to our Bioanalyzer, RNA-seq, and RT-qPCR analyses that our three-stage purification pipeline produces highly pure mRNA. Commonly, mRNA modification LC-MS/MS analyses characterize only a select few target modifications, which prevents the utilization of LC-MS/MS to judge purity of mRNA. The LC-MS/MS assay described here quantifies up to 46 ribonucleoside modifications in a single analysis, enabling us to use our method to thoroughly characterize mRNA purity. Our analyses ensure that rRNA and tRNA specific modifications are not present at a detectable level in our highly purified mRNA. This highly sensitive corroboration of our Bioanalyzer findings is essential because RNA-seq is not able to sufficiently report on tRNA contamination without utilization of more tRNA compatible reverse transcriptases.

**Figure 4:**
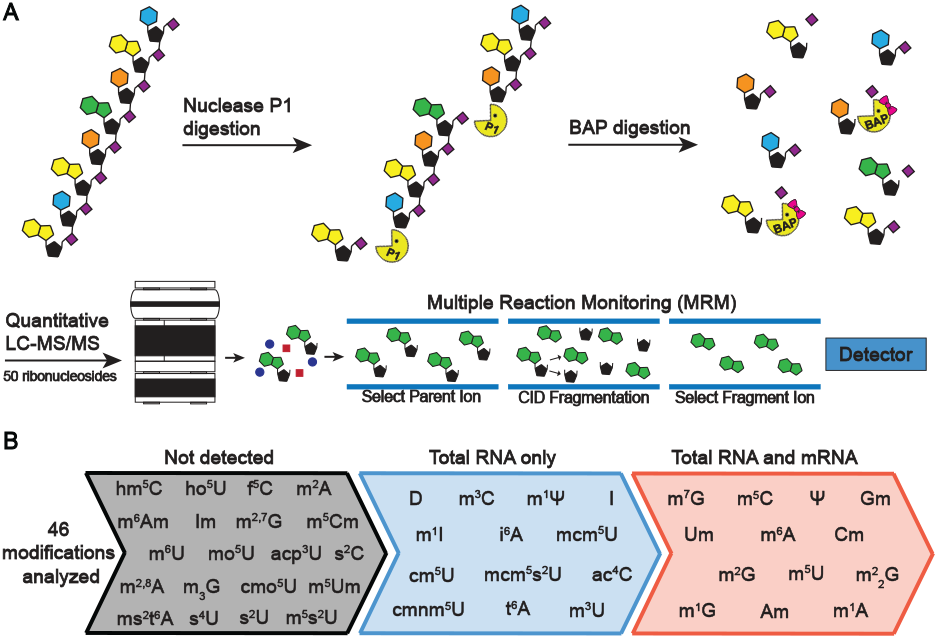
Enzymatic digestion and LC-MS/MS analysis of *S. cerevisiae* total RNA and mRNA. **A)** RNA is enzymatic digested to ribonucleosides through a two-stage process. RNA is first digested to nucleotide monophosphates by nuclease P1 and then dephosphorylated to ribonucleosides by bacterial alkaline phosphatase. The resulting ribonucleosides are separated using reverse phase chromatography and then quantified using MRM on a triple quadrupole mass spectrometer. **B)** *S. cerevisiae* total RNA and mRNA were analyzed using the LC-MS/MS method developed to quantify 46 modifications in a single analysis. In total RNA, 26 modifications were detected while 13 ribonucleosides were detected in the highly purified mRNA.

Since all RNA present in our samples will be enzymatically degraded to ribonucleosides during sample preparation **(Figure 4A)**, contaminating highly modified ncRNA will lead to inaccurate modifications quantification in mRNA samples. Thus, extensive quality control for mRNA purity is necessary to give us confidence in downstream LC-MS/MS analyses; however, such data are rarely provided in previous mRNA modification LC-MS/MS studies. Together, we provide four types of evidence (Bioanalyzer, RT-qPCR, RNA-seq, and LC-MS/MS) that our protocol yields highly pure mRNA appropriate for LC-MS/MS analysis. The incorporation of tRNA compatible reverse transcriptases into our RNA-seq pipeline would further confirm the removal of tRNA contamination in addition to Bioanalyzer and LC-MS/MS. While previous mRNA purification pipelines may inaccurately portray the modification landscape, this pipeline will enable the accurate characterization and quantification of mRNA modifications by providing highly purified mRNA for the analysis using solely commercial kits. We believe that our purification and rigorous purity assessment pipeline could provide a standard method to purify polyadenylated mRNA from total RNA for LC-MS/MS analysis.

### m^1^G, m^2^G, m^2^_2_G, and m^5^U detected in *S. cerevisiae* mRNA

In our purified mRNA samples, we detected 13 ribonucleoside modifications that ranged in abundance from pseudouridine (0.023 Ψ/U%) to 1-methyladenosine (0.00014 m^1^A/A%) **(Figure 4B, Supplemental Figure S6 and Supplemental Tables S5 and S6)**. These abundances are lower than other previous mRNA modification LC-MS/MS analyses, including a previous *S. cerevisiae* study^20^. We attribute this to the fact that our mRNA is more pure than the mRNA used in previous studies, which leads to lower modification abundances in our samples since there is less contaminating highly modified ncRNA. Most of these modifications we observed in our samples are known to be present in *S. cerevisiae* mRNA; however, we detected four modifications for the first time in *S. cerevisiae* (m^1^G, m^2^G, m^2^_2_G, and m^5^U) **(Figure 5A)**. This finding corroborates previous studies that detected m^1^G^24^ and m^5^U^15,21,91^ in *Arabidopsis thaliana* and multiple mammalian cell lines at similar levels, respectively.

**Figure 5:**
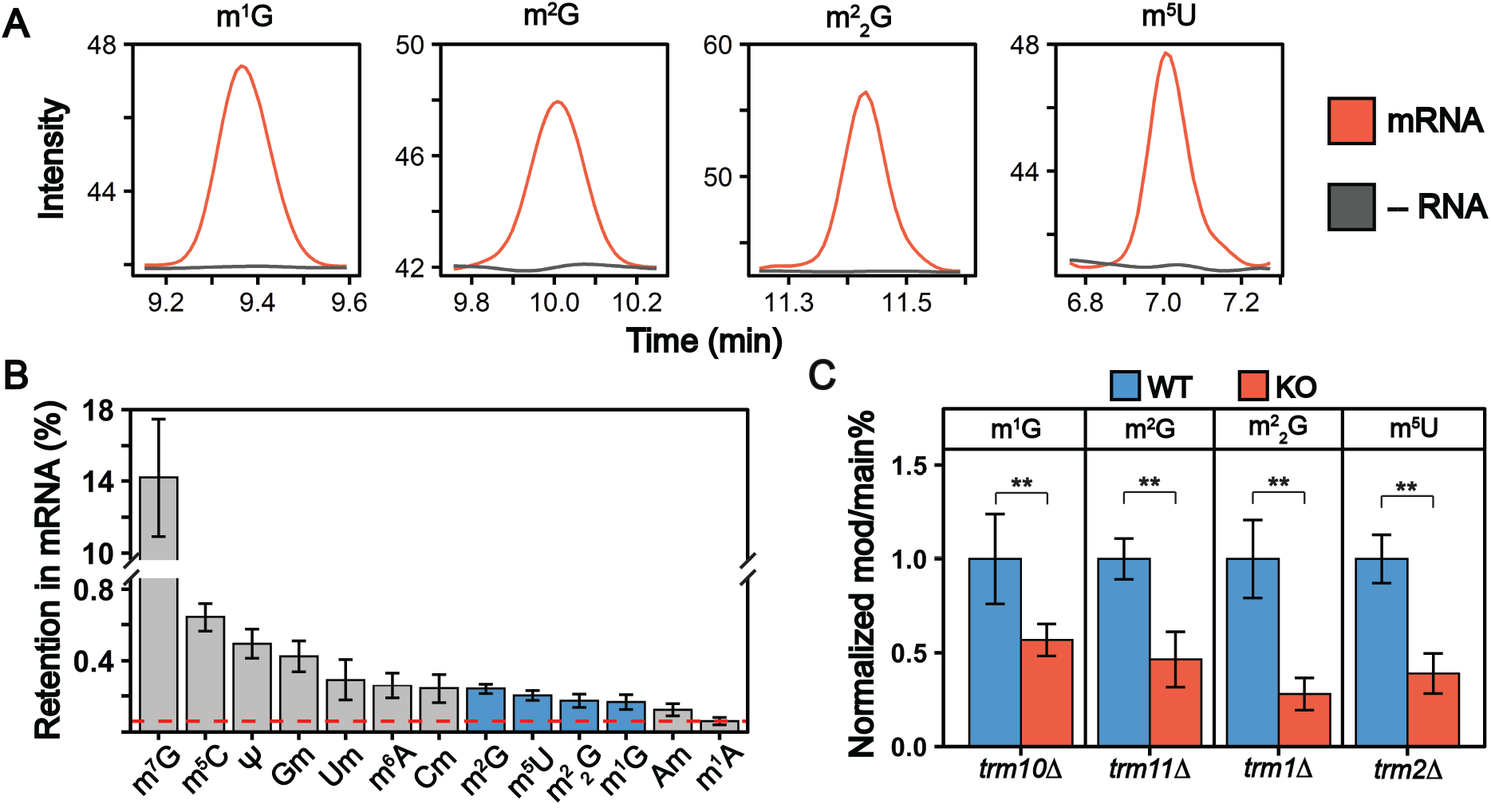
m^1^G, m^2^G, m^2^ G, and m^5^U are present in *S. cerevisiae* mRNA. **A)** Overlaid extracted ion chromatograms displaying m^1^G, m^2^G, m^2^ G, and m^5^U are detected in our mRNA samples (red) above our digestion control samples without RNA added (grey). **B)** m^1^G, m^2^G, m^2^ G, and m^5^U are only present in *S. cerevisiae* tRNA; thus, we reasoned that they would be retained at a higher percentage than other highly abundant tRNA modifications if they are present in mRNA. Dihydrouridine, which is the most abundant non-mRNA modification in tRNA, was not detected in our purified mRNA samples. If dihydrouridine existed at levels just below our limit of detection (530 amol), the maximum retention of solely tRNA modifications would be 0.06% (red dashed line). The four new mRNA modifications we detect, along with all other known mRNA modifications, are retained at greater extents which proves these modifications exist in *S. cerevisiae* mRNA. The error bars are the standard deviation of the percent retention. **C)** m^1^G, m^2^G, m^2^ G, and m^5^U are incorporated into *S. cerevisiae* mRNA by their corresponding tRNA modifying enzymes (Trm10, trm11, Trm1, and Trm2 respectively). The modification/main base% (e.g., m^1^G/G%) were normalized to their levels in the average WT mRNA levels. A significant decrease (**p < 0.01) was detected for all cases. The error bars are the standard deviation of the normalized mod/main base%.

We next critically considered our findings and contemplated the possibility that the signals we detect originated from minor contaminations of tRNA. Prior to this study, in *S. cerevisiae* m^1^G, m^2^G, m^2^_2_G, and m^5^U have only been reported in tRNA^3,92^. Therefore, we reasoned that if these methylated nucleosides are present in *S. cerevisiae* mRNA, they must be retained at higher levels than other tRNA modifications that are not found in mRNA. DHU is the second most abundant RNA modification in *S. cerevisiae* tRNA and thus provides a measure of maximum tRNA contamination **(Supplemental Table S6)**. We did not detect any DHU in our purified mRNA samples. Recent sequencing based studies have reported the presence of DHU in mammalian and *S. pombe* mRNA^15,16^, but our findings indicate DHU either does not exist within *S. cerevisiae* mRNA or is incorporated at levels below our limit of detection. If dihydrouridine existed at levels just below our limit of detection (530 amol), **(Supplemental Table S1)** the maximum extent of DHU/U% retention in our purified mRNA would be 0.06% when calculated using the average digest uridine concentration in a sample of digested mRNA. We find that m^1^G, m^2^G, m^2^_2_G and m^5^U (in addition to all other modifications) were retained to a greater extent than the maximum theoretical retention of level of DHU (>2.5-fold more) in our purified mRNA **(Figure 5B and Supplemental Table S7)**.

Since all contaminating ncRNA species will be digested to ribonucleosides along with mRNA, it is essential to carefully assess our mRNA purity quality controls and the retention of known exclusive ncRNA modifications in our mRNA modification LC-MS/MS data. In this work, our extensive mRNA purity quality control by Bioanalyzer, RNA-seq, RT-qPCR, and LC-MS/MS in conjunction with there being no other exclusive highly abundant tRNA and rRNA modifications detected in our purified mRNA samples confirms that these modifications are present in *S. cerevisiae* mRNA.

### Trm1, Trm2, Trm10 and Trm11 incorporate methylated guanosine and uridine modifications into *S. cerevisiae* mRNA

Many of the reported mRNA modifications are incorporated by the same enzymes that catalyze their addition into tRNAs and rRNAs^3^. We investigated if the enzymes responsible for inserting m^1^G, m^2^G, m^2^_2_G, and m^5^U into *S. cerevisiae* tRNAs (Trm10, Trm11, Trm1, and Trm2 respectively) also incorporate them into *S. cerevisiae* mRNA. We compared the levels of m^1^G, m^2^G, m^2^_2_G, and m^5^U in mRNA purified from wild-type and mutant (*trm10*Δ, *trm11*Δ, *trm1*Δ, and *trm2*Δ) *S. cerevisiae*. The abundance of all four modifications decreased significantly in mRNAs purified from the knockout cell lines **(Figure 5C and Supplemental Tables S6)**. While this demonstrates that the tRNA modifying enzymes incorporate these modifications into *S. cerevisiae* mRNA, low levels of m^1^G, m^2^G, m^2^_2_G, and m^5^U modifications are still detected in the mRNAs from knockout cell lines **(Figure 5C)**. Several explanations could account for this. A second enzyme, Trm5, also catalyzes m^1^G addition into tRNAs and could possibly explain the remaining mRNA m^1^G signals. However, given that m^1^G and m^2^G were previously found as minor products of methylation damage in DNA and RNA^31,93–99^, it is perhaps more likely that the remaining low-level signals that we detect arise from methylation associated RNA damage or minor off target methylation by other enzymes. Regardless of how they are incorporated, when present, these modifications have the potential to impact mRNA function.

### m^1^G, m^2^G and m^5^U containing mRNA codons slow amino acid addition by the ribosome in a position dependent manner

While our LC-MS/MS assays indicate that m^1^G, m^2^G, m^2^_2_G and m^5^U modifications exist within *S. cerevisiae* mRNA, no previous work has revealed the location or biological consequence of these modifications in mRNA. Despite their low abundance compared to ncRNA modifications (typically significantly lower than 1% modified), evidence that mRNA modifications can alter the chemical and topological properties of modified transcripts which resultingly affect their stability and function continues to increase. Analogously, N-linked and O-linked glycosylations of proteins occur at rates less than approximately 1% and 0.04% per target amino acid, respectively^100^; however, these post-translational modifications play important biological roles, such protein localization and receptor interaction^101,102^, and their misregulation is linked to multiple diseases^103^ despite their low abundance. mRNAs are all substrates for the ribosome, and post-transcriptional modifications can change how the ribosome decodes a message by altering the hydrogen bonding patterns between the mRNA codons and aminoacylated-tRNAs^104–109^. Indeed, several mRNA modifications have been shown to alter the overall rate and fidelity of protein synthesis in a modification and codon-position dependent manner^40,41,110–115^. Such perturbations to protein synthesis can have significant consequences even when modifications are incorporated into mRNAs transcripts at very low levels, as exemplified by the biological consequences of oxidatively damaged mRNAs, which exist at levels similar to m^1^G, m^2^G, m^2^_2_G and m^5^U^31,116^. We investigated how the insertion of m^5^U, m^1^G, and m^2^G into mRNA codons impacts translation using a well-established reconstituted *in vitro* translation system^40^ **(Figure 6A)**. This system has long been used to investigate how the ribosome decodes mRNAs because it can be purified in sufficient quantities to conduct high-resolution kinetic studies. Translation elongation is well conserved between bacteria and eukaryotes^117^, and prior studies demonstrate that mRNA modifications (e.g. pseudouridine, N6-methyladenosine and 8-oxo-G) that slow elongation and/or change mRNA decoding elongation in the reconstituted *E. coli* system^40,41,110,118^ also do so in eukaryotes^40,119–121^. m^2^_2_G was not selected for study because the phosphoramidite required for mRNA oligonucleotide synthesis is not commercially available.

**Figure 6:**
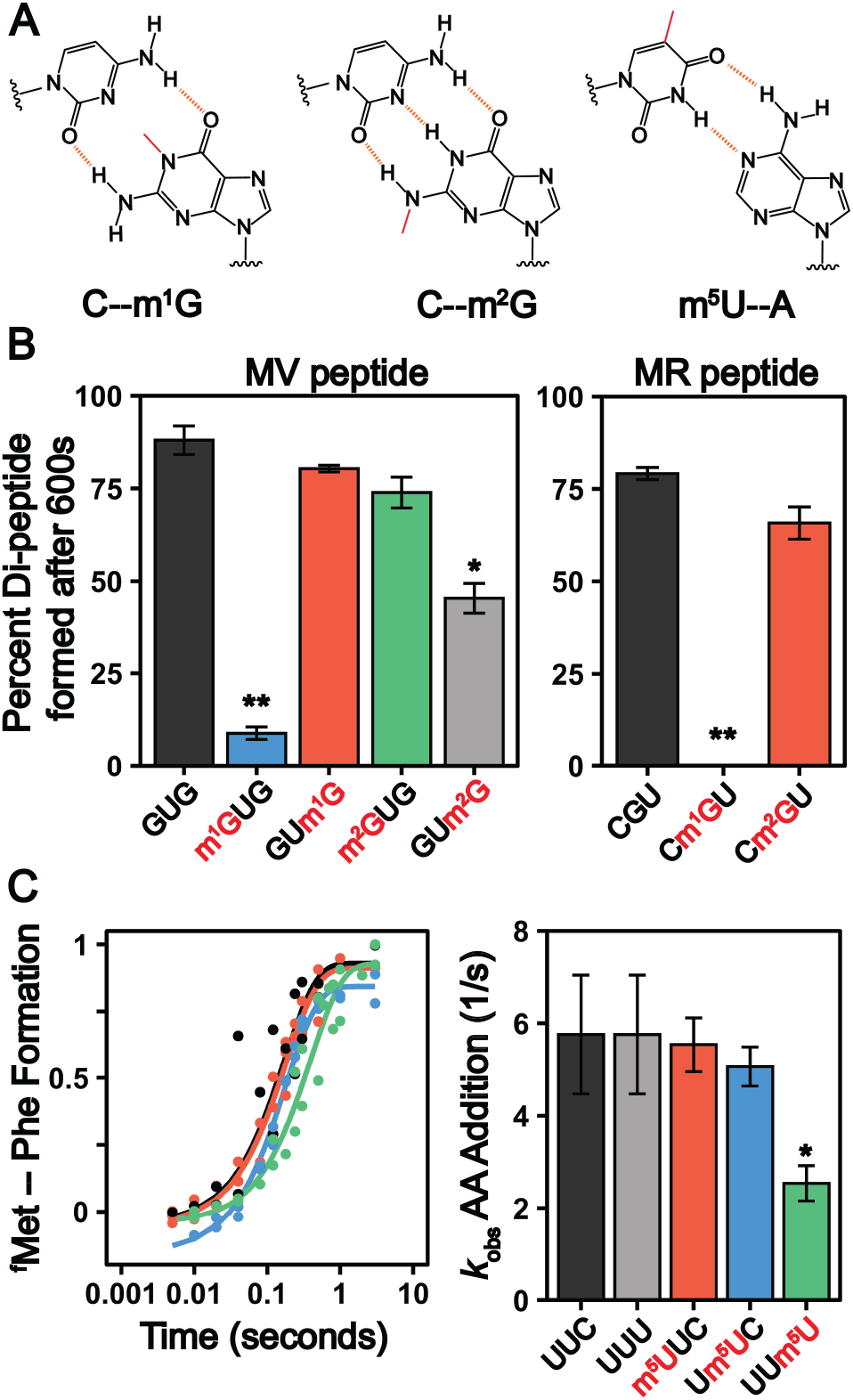
Methylated guanosine and uridine modifications alter amino acid addition. **A)** Watson-Crick base pairing of m^1^G, m^2^G and m^5^U. The added methylation is displayed in red and the hydrogen bond interactions displayed as a dashed orange line. **B)** Total peptide formation of translation reactions after 600 seconds using transcribed or single-nucleotide modified mRNAs encoding for either (Left Panel) Met-Val (GUG) or (Right Panel) Met-Arg (CGU) dipeptide. Error bars are the standard deviation. **B)** Time courses displaying the formation of ^f^Met-Phe dipeptide on an unmodified and singly modified UUC or UUU codons (left panel). Observed rate constants (right panel) were determined from the fit data. The error bars are the standard deviation of the fitted value of *k*_obs_.

In our assays, 70S ribosome initiation complexes (ICs) containing ^35^S-^f^Met-tRNA^fmet^ programmed in the A site are formed on transcripts encoding Met-Phe, Met-Arg, or Met-Val dipeptides. Ternary complexes comprised of aminoacyl-tRNA:EF-Tu:GTP are added to the ICs to begin translation. Reactions are quenched as desired timepoints by KOH, and the unreacted ^35^S-^f^Met-tRNA^fmet^ and dipeptide translation products are visualized by electrophoretic TLC (eTLC) **(Supplementary Figures S7 through S10)**. We evaluated the extent of total dipeptide synthesis and/or the rate constants (*k*_*obs*_) for amino acid incorporation on unmodified (CGU, GUG, UUC, UUU) and modified (Cm^1^GU, Cm^2^GU, m^1^GUG, m^2^GUG, GUm^1^G, GUm^2^G, m^5^UUC, Um^5^UC, Uum^5^U) codons. The presence of modifications in the codons were verified by direct infusion ESI-MS or nanoelectrospray ionization (nESI)-MS **(Supplemental Figures S11 to S13)**. We observed that the extent of amino acid addition is drastically reduced when m^1^G is present at the first or second position in a codon but is restored to normal levels when m^1^G is at the third nucleotide **(Figure 6B and Supplemental Figures S7 through S9)**. Codons containing m^2^G show a more modest defect in dipeptide production, only significantly impeding dipeptide synthesis (1.9 ± 0.2-fold) when m^2^G is in the third position of a codon **(Figure 6B and Supplemental Figures S7 through S9)**. These findings are consistent with a previous report indicating that insertion of a single m^1^G and m^2^G modification into an mRNA codon reduces the overall protein production and translation fidelity in a position and codon dependent manner^115^. m^1^G and m^2^G should both disrupt Watson-Crick base pairing between mRNAs and tRNAs **(Figure 6A)** and might be expected to alter amino acid addition in similar ways. However, our results reveal that the insertion of m^1^G has a much larger consequence than m^2^G on peptide production. This can be partially rationalized by the fact that m^1^G would impede canonical Watson-Crick base-pairing by eliminating a central H-bond interaction, while m^2^G disrupts only peripheral interactions **(Figure 6A)**. Additionally, the methylation of the analogous position of adenosine (m^1^A) similarly abolishes the ability of the ribosome to add amino acids^30^, suggesting that the conserved N1 position on purine nucleobases is particularly crucial to tRNA decoding. The hydrogen bonding patterns possible between m^2^G and other nucleosides would be expected to closely resemble those of another well studied modification, inosine. Inosine also has a moderate (if any) impact on the rates of protein synthesis, though it can promote amino acid mis-incorporation^122,123^. The limited consequence of both inosine and m^2^G on overall peptide production indicates that purine peripheral amines on the Watson-Crick face are less important than the N1 position for ensuring the rapid addition of amino acids by the ribosome.

In contrast to the guanosine modifications that we investigated, transcripts containing m^5^U Phe-encoding codons did not reduce the total amount of dipeptide produced **(Figure 6C)**. However, the insertion of m^5^U into codons can reduce the rate constants for amino acid addition (*k*_obs_) in a position dependent manner, similar to Ψ modified transcripts^40^. The rate constant for Phe incorporation on an unmodified and modified codons at the 1^st^ and 2^nd^ position were comparable to an unmodified codon, with a *k*_obs_ of ∼ 5s^-1^ **(Figure 6C and Supplemental Figure S10)**. However, when m^5^U is in the 3^rd^ position we see a 2-fold decrease in the *k*_obs_ at ∼ 2.5s^-1^ **(Figure 6C and Supplemental Figure S10)**. This is the first evidence that m^5^U can influence amino acid addition when encountered by the ribosome. It is less clear how m^5^U and other modifications that do not change the Watson-Crick face of nucleobases (e.g., Ψ and 8-oxoG) impact translation^124^. It is possible that such modifications alter nucleobase ring electronics to perturb the strength of the hydrogen bond donors and acceptors involved in base pairing.

While the levels of the mRNA modifications we identified are lower than that of more well-established modifications (m^6^A and Ψ), our findings suggest that they still have potential to impact biology. Although our data do not report on the ability of the modifications that we uncover to control gene expression or identify the number of mRNAs that they are in, they do suggest that there will be consequences for translation when these modifications are encountered by the ribosome. It is also important to note that the levels and distributions of mRNA modifications (enzymatic and RNA damage) can change significantly in response to different environmental conditions, so the low levels of modification that we measure in healthy, rapidly growing yeast have the potential to significantly increase under stress^20,28,116,125^. The three modifications we investigated alter translation differently depending on their location within a codon. Such a context dependence has been observed for every mRNA modification investigated to date^124^. Modifications have the capacity to change intra-molecular interactions with an mRNA, or interactions between rRNA and mRNA within the A site. There is growing evidence that such factors, and not only anticodon:codon interactions, have a larger contribution to translation elongation than previously recognized. For example, ribosome stalling induced by the rare 8-oxo-guanosine damage modification has the potential to perturb ribosome homeostasis or even the small pauses in elongation induced by mRNA pseudouridine modifications can impact levels of protein expression in a gene specific manner^31,121^. Additionally, transient ribosome pauses have the potential modulate co-translational protein folding or provide time for RNA binding proteins to interact with a transcript^126,127^. Future systematic biochemical and computational studies are needed to uncover the causes of the context dependence. Additionally, the continued development of RNA-seq technologies is needed to locate these modifications throughout the transcriptome. This information will be broadly useful as researchers seek to identify which of the modified mRNA codons are the most likely to have molecular level consequences when encountered by a translating ribosome.

## CONCLUSIONS

Mass spectrometry based approaches are widely used to study protein post-translational modifications, but the application of similar techniques to investigate mRNA post-transcriptional modifications has not been widely adopted. The current LC-MS/MS workflows for discovering and studying mRNA modifications are hindered by either low-throughput method development, inadequate mRNA purification, or insufficient sensitivities to detect low level mRNA modifications. This study presents mRNA purification, validation, and LC-MS/MS pipelines that enable the sensitive and highly multiplexed analysis of mRNA and ncRNA modifications. These developments enable us to confidently identify four previously unreported mRNA modifications in *S. cerevisiae* (m^1^G, m^2^G, m^2^_2_G and m^5^U), demonstrating the utility of applying LC-MS/MS to discover and quantify mRNA modifications. In addition to revealing the enzymes that incorporate these modifications, we also demonstrate that the presence of m^1^G, m^2^G, and m^5^U in mRNA can impede translation. However, the impacts of the modifications on amino acid addition are not uniform, with the position and identity of each modification resulting in a different outcome on dipeptide production. This work suggests that the ribosome will regularly encounter a variety of modified codons in the cell and that depending on the identity and position of the modification, these interactions can alter the elongation step in protein synthesis.

## METHODS

### *S. cerevisiae* cell growth and mRNA purification

Wild-type, *Δtrm1, Δtrm2, Δtrm10* and *Δtrm11* BY4741 *S. cerevisiae* (Horizon Discovery) were grown in YPD medium as previously described^20^. Knockout cells lines were grown with 200 μg/mL Geneticin. Briefly, 100 mL of YPD medium was inoculated with a single colony selected from a plate and allowed to grow overnight at 30°C and 250 RPM. The cells were diluted to an OD_600_ of 0.1 with 300 mL of YPD medium and were grown to an OD_600_ of 0.6-0.8 at 30°C and 250 RPM. The cell suspension was pelleted at 3,220 x g at 4°C and used for the RNA extraction.

*S. cerevisiae* cells were lysed as previously described with minor alterations^20,128^. The 300 mL cell pellet was resuspended in 12 mL of lysis buffer (60 mM sodium acetate pH 5.5, 8.4 mM EDTA) and 1.2 mL of 10% SDS. One volume (13.2 mL) of acid phenol:chloroform:isoamyl alcohol (125:24:1; Sigma-Aldrich, USA; P1944) was added and vigorously vortexed. The mixture was incubated in a water bath at 65°C for five min and was again vigorously vortexed. The incubation at 65°C and vortexing was repeated once. Then, the mixture was rapidly chilled in an ethanol/dry ice bath until lysate was partially frozen. The lysate was allowed to thaw and then centrifuged for 15 min at 15,000 x g. The upper layer containing the total RNA was washed three additional times with 13.2 mL phenol and the phenol was removed using two chloroform extractions of the same volume. The resulting RNA was ethanol precipitated in the presence of 1/10^th^ volume of 3 M sodium acetate and then a second time in the presence of 1/2 volume of 7.5 M ammonium acetate. The extracted total RNA was treated with 140 U RNase-free DNase I (Roche, 10 U/μL) in the supplied digestion buffer at 37°C for 30 min. The DNase I was removed through an acid phenol-chloroform extraction. The resulting RNA was ethanol precipitated in the presence of 1/10^th^ volume of 3 M sodium acetate pH 5.2 and then a second time in the presence of 1/2 volume of 7.5 M ammonium acetate. The precipitated RNA was pelleted and resuspended in water. The resulting total RNA was used for our LC-MS/MS, bioanalyzer, and RNA-seq analyses.

mRNA was purified through a three-stage purification pipeline. First, small RNA (tRNA and 5S rRNA) was diminished from 240 μg of total RNA using a Zymo RNA Clean and Concentrator-100 kit to purify RNA > 200nt. Two consecutive poly(A) enrichment steps were applied to 125 μg of the resultant small RNA diminished samples using Dynabeads oligo-dT magnetic beads (Invitrogen, USA). The resulting poly(A) RNA was ethanol precipitated using 1/10^th^ volume of 3 M sodium acetate pH 5.2 and resuspended in 14 μL of water. Then, we removed the residual 5S, 5.8S, 18S, and 28S rRNA using the commercial riboPOOL rRNA depletion kit (siTOOLs Biotech). The Bioanalyzer RNA 6000 Pico Kit (Agilent) was used to evaluate the purity of the mRNA prior to LC-MS/MS analysis.

### qRT-PCR

DNase I treated total RNA and three-stage purified mRNA (200 ng) were reverse transcribed using the RevertAid First Strand cDNA Synthesis Kit (Thermo Scientific) using the random hexamer primer. The resulting cDNA was diluted 5000-fold and 1 μL of the resulting mixture was analyzed using the Luminaris Color HiGreen qPCR Master Mix (Thermo Scientific) with gene-specific primers (**Supplemental Table S8**).

### RNA-seq

The WT *S. cerevisiae* mRNA was analyzed by RNA-seq as previously described with minimal alterations^20^. Briefly, 50 ng of DNase I treated total RNA and three-stage purified mRNA from the two biological replicates were fragmented using the TruSeq RNA Library Prep Kit v2 fragmentation buffer (Illumina). First-strand cDNA synthesis was performed using the random hexamer primer, and the second strand was synthesized using the Second Strand Master Mix. The resulting cDNA was purified with AMPure XP beads (Beckman Coulter), the ends were repaired, and the 3’ end was adenylated. Lastly, indexed adapters were ligated to the DNA fragments and amplified using 15 PCR cycles. Paired-end sequencing was performed for the cDNA libraries using 2.5% of an Illumina NovaSeq (S4) 300 cycle sequencing platform flow cell (0.625% of flow cell for each sample). All sequence data are paired-end 150 bp reads.

FastQC (v0.11.9)^129^ was used to evaluate the quality of the raw and trimmed reads. Then, cutadapt (v1.18)^130^ was used to trim to paired-end 50 bp reads and obtain high quality clean reads with the arguments -u 10 -U 10 -l 50 -m 15 -q 10. Following, Bowtie2 (v2.2.5)^131^ was used to align the forward strand reads to *S. cerevisiae* reference genome (R64-1-1) with the default parameters. Following alignment, Rmmquant tool R package (v1.6.0)^132^ and the gene_biotype feature in the *S. cerevisiae* GTF file was used to count the number of mapped reads for each transcript and classify the RNA species, respectively.

### RNA digestions and LC-MS/MS analysis

RNA (200 ng) was hydrolyzed to composite mononucleosides using a two-step enzymatic digestion. The RNA was first hydrolyzed overnight to nucleotide monophosphates using 300 U/ μg Nuclease P1 (NEB, 100,000 U/mL) at 37°C in 100 mM ammonium acetate (pH 5.5) and 100 μM ZnSO_4_. Following, the nucleotides were dephosphorylated using 50 U/μg bacterial alkaline phosphatase (BAP, Invitrogen, 150U/μL) for 5 hrs at 37°C in 100 mM ammonium bicarbonate (pH 8.1) and 100 μM ZnSO_4_. Prior to each reaction, the enzymes were buffer exchanged into their respective reaction buffers above using a Micro Bio-Spin 6 size exclusion spin column (Biorad) to remove glycerol and other ion suppressing constituents. After the reactions, the samples were lyophilized and resuspended in 9 μL of water and 1 μL of 400 nM ^15^N_4_-inosine internal standard.

The resulting ribonucleosides were separated using a Waters Acquity HSS T3 column (1 × 100 mm, 1.8 μm, 100 Å) with a guard column at 100 μL/min on a Agilent 1290 Infinity II liquid chromatograph interfaced to a Agilent 6410 triple quadrupole mass spectrometer. Mobile phase A was 0.01% (v/v) formic acid in water and mobile phase B was 0.01% (v/v) formic acid in acetonitrile. The gradient is displayed in **Supplemental Table S9**. The autosampler was held at 4°C, and 5 μL was injected for each sample. The eluting ribonucleosides were quantified using MRM and ionized using electrospray ionization in positive mode at 4 kV **(Supplemental Table S10)**. The electrospray ionization conditions were optimized by infusing 500 nM uridine at 100 μL/min at 5% mobile phase B. The gas temperature was 350°C, the gas flow rate was 10 L/min, and the nebulizer gas pressure was 25 psi. After each RNA digestion sample, a wash gradient injection was performed to eliminate any column carryover of late eluting nucleosides (e.g., i^6^A) **(Supplemental Table S9)**.

To compare the sensitivity between the 1 mm and 2 mm I.D. column chromatographies, a 2.1 mM Waters Acquity HSS T3 column (2.1 × 100 mm, 1.8 μm, 100A) with a guard column was used at 400 uL/min using the same gradient and mobile phases described above. The source conditions for the 2.1 mm I.D. column were optimized by infusing 500 nM uridine at 400 μL/min at 5% mobile phase B. The gas temperature was 350°C, the gas flow rate was 10 L/min, and the nebulizer gas pressure was 55 psi. For both analyses, 5 uL of ribonucleoside standard mixes containing 1.4 μM canonical nucleosides and 72 nM modifications was injected.

To quantify RNA nucleosides calibration curves were created for the four main bases, 45 natural modified nucleosides, and 1 non-natural modified nucleoside using seven calibration points ranging over four orders of magnitude. ^15^N_4_-inosine (40 nM) was used as the internal standard for all ribonucleosides. The concentrations of ribonculeoside in the calibration curves standards can be found in **Supplemental Table 11**. Suppliers for ribonucleoside standards can be found in **Supplemental Table 12**. Automated peak integration was performed using the Agilent MassHunter Workstation Quantitative Analysis Software. All peaks were visually inspected to ensure proper integration. The calibration curves were plotted as the log_10_(response ratio) versus the log_10_(concentration (pM)) and the RNA sample nucleoside levels were quantified using the resulting linear regression. The limits of detection were calculated using:

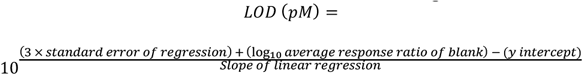

The calculated LOD was then converted to amol. For each RNA enzymatic digestion samples, the respective calibration curve was used to calculate nucleoside concentrations in the samples.

The retention of modifications in mRNA was calculated using the following equation:

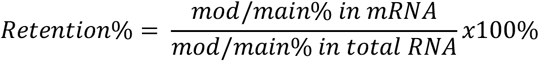

### *E. coli* ribosomes and translation factor purification

Ribosomes were purified from *E. coli* MRE600 as previously described^40^. All constructs for translation factors were provided by the Green lab unless specifically stated otherwise. The expression and purification of translation factors were carried out as previously described^40^.

### Preparation of tRNA and mRNA for *in vitro* translation assays

Unmodified transcripts were prepared using run-off T7 transcription of Ultramer DNA templates that were purchased from Integrated DNA Technologies **(Supplemental Table S13)**. HPLC purified modified mRNA transcripts containing 5-methyluridine, 1-methylguanosine, and N2-methylguanosine were purchased from Dharmacon **(Supplemental Table S14)**. The homogeneity and accurate mass for most of the purchased modified oligonucleotides were confirmed by direct infusion ESI-MS prior to use by Dharmacon **(Supplementary Figure S11 through S13)**. For the remaining purchased oligonucleotides lacking Dharmacon spectra, they were analyzed on a ThermoFisher Q-Exactive UHMR Hybrid Quadrupole-Orbitrap Mass Spectrometer in a negative ionization polarity. Samples were buffer exchanged into 100 mM ammonium acetate (AmOAc) using Micro Bio-Spin P-6 gel columns and directly infused via nanoelectrospray ionization (nESI). nESI was performed using borosilicate needles pulled and coated in-house with a Sutter p-97 Needle Puller and a Quorum SCX7620 mini sputter coater, respectively. The acquired native mass spectra were deconvoluted using UniDec^133^ in negative polarity **(Supplementary Figure S11)**.

Native tRNA was purified as previously described with minor alterations^134^. Bulk *E. coli* tRNA was either bought in bulk from Sigma-Aldrich or purified from a HB101 *E. coli* strain containing pUC57-tRNA that we obtained from Prof. Yury Polikanov (University of Illinois, Chicago). Two liters of media containing Terrific Broth (TB) media (TB, 4 mL glycerol/L, 50 mM NH_4_Cl, 2 mM MgSO_4_, 0.1 mM FeCl_3_, 0.05% glucose and 0.2% lactose (if autoinduction media was used)) were inoculated with 1:400 dilution of a saturated overnight culture and incubated with shaking at 37°C overnight with 400 mg/ml of ampicillin. Cells were harvested the next morning by 30 min centrifugation at 5000 RPM and then stored at -80°C. Extraction of tRNA was done by first resuspending the cell pellet in 200 mL of resuspension buffer (20 mM Tris-Cl, 20 mM Mg(OAc)_2_ pH 7.) The resuspended cells were then placed in Teflon centrifuge tubes with ETFE o-rings containing 100 mL acid phenol/chloroform/isoamyl alcohol mixture. The tubes were placed in a 4°C incubator and left to shake for 1 hr. After incubation, the lysate was centrifuged for 60 min at 3,220 x g at 4°C. The supernatant was transferred to another container and the first organic phase was then back-extracted with 100mL resuspension buffer and centrifuged down for 60 min at 3,220 x g at 4°C. Aqueous solutions were then combined and a 1/10 volume of 3 M sodium acetate pH 5.2 was added and mixed well. Isopropanol was added to 20% and after proper mixing was centrifuged to remove DNA at 13,700 x g for 60 min at 4°C. The supernatant was collected, and isopropanol was added to 60% and was left to precipitate at -20°C overnight. The precipitated RNA was pelleted at 13,700 x g for 60 min at 4°C and resuspended with approximately 10 mL 200 mM Tris-Acetate, pH 8.0. The RNA was incubated at 37°C for at least 30 min to deacylate the tRNA. After incubation 1/10^th^ volume of 3 M sodium acetate pH 5.2 and 2.5 volumes of ethanol was added to precipitate the RNA. Then, the mixture was centrifuged at 16,000 x g for 60 min at 4°C. The pellet was washed with 70% ethanol, resuspended in water, and desalted using an Amicon 10 kDa MWCO centrifugal filter prior to purification (Millipore-Sigma, USA).

Next, the tRNA was isolated using a Cytiva Resource Q column (6 mL) on a AKTA Pure 25M FPLC. Mobile phase A was 50 mM NH_4_OAc, 300 mM NaCl, and 10 mM MgCl_2_. Mobile phase B was 50 mM NH_4_OAc, 800 mM NaCl, 10 mM MgCl_2_. The resuspended RNA was filtered, loaded on the Resource Q column, and eluted with a linear gradient from 0-100% mobile phase B over 18 column volumes. Fractions were pulled and ethanol precipitated overnight at - 20°C.

The precipitated RNA was resuspended in water and filtered prior to purification on a Waters XBridge BEH C18 OBD Prep wide pore column (10 × 250 mm, 5 μm). Mobile phase A was 20 mM NH_4_OAc, 10 mM MgCl_2_, and 400 mM NaCl at pH 5 in 100% water. Mobile phase B was 20 mM NH_4_OAc, 10 mM MgCl_2_, and 400 mM NaCl at pH 5 in 60% methanol. The injection volume was 400 μl. A linear gradient of mobile phase B from 0-35% was done over 35 min. After 35 min, the gradient was increased to 100% mobile phase B over 5 min and held at 100% for 10 min, column was then equilibrated for 10 column volumes before next injection with mobile phase A. TCA precipitations were performed on the fractions to identify fractions containing the phenylalanine tRNA as well as measuring the A_260_ and amino acid acceptor activity.

### Formation of *E. coli* ribosome initiation complexes

Initiation complexes (ICs) were formed in 1X 219-Tris buffer (50 mM Tris pH 7.5, 70 mM NH_4_Cl, 30 mM KCl, 7 mM MgCl_2_, 5 mM β-ME) with 1 mM GTP as previously described^134^. 70S ribosomes were incubated with 1 μM mRNA (with or without modification), initiation factors (1, 2, and 3) all at 2 μM final, and 2 μM of radiolabeled ^35^S- ^f^Met-tRNA^fMet^ for 30 min at 37°C. After incubation, MgCl_2_ was added to a final concentration of 12 mM. The ribosome mixture was then layered onto 1 mL cold buffer D (20 mM Tris-Cl, 1.1 M sucrose, 500 mM NH_4_Cl, 10 mM MgCl_2_, 0.5 mM disodium EDTA, pH 7.5) and centrifuged at 69,000 rpm for 2 hrs at 4°C. After pelleting, the supernatant was discarded into radioactive waste, and the pellet was resuspended in 1X 219-tris buffer and stored at -80°C.

### *In vitro* amino acid addition assays

IC complexes were diluted to 140 nM with 1X 219-Tris buffer. Ternary complexes (TCs) were formed by first incubating the EF-Tu pre-loaded with GTP (1X 219-Tris buffer, 10 mM GTP, 60 μM EFTu, 1 μM EFTs) at 37°C for 10 min. The EF-Tu mixture was incubated with the tRNA mixture (1X 219-Tris buffer, Phe-tRNA^Phe^ (1-10 μM), 1 mM GTP) for another 15 min at 37°C. After TC formation was complete, equal volumes of IC complexes (70 nM) and ternary complex (1 μM) were mixed either by hand or using a KinTek quench-flow apparatus. Discrete time-points (0-600 seconds) were taken as to obtain observed rate constants on m^5^U-containing mRNAs. Each time point was quenched with 500 mM KOH (final concentration). Time points were then separated by electrophoretic TLC and visualized using phosphorescence as previously described^40,134^. Images were quantified with ImageQuant. The data were fit using Equation 1:

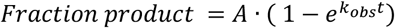

## Supporting information

Jones_etal_Supplemental Figures

Jones_etal_Supplemental Tables

## ASSOCIATED CONTENT

### Supporting Information

Supporting Figures and Supporting Tables files are available.

### Author Contributions

<u>Joshua D. Jones</u>: Conceptualization, Methodology, Investigation, Formal Analysis, Writing – Original Draft, Writing

– Review & Editing, Visualization

<u>Monika K. Franco</u>: Conceptualization, Investigation, Formal Analysis, Writing – Original Draft, Writing – Review & Editing

<u>Tyler J. Smith</u>: Conceptualization, Investigation, Formal Analysis, Writing – Original Draft, Writing – Review & Editing

<u>Laura R. Snyder</u>: Investigation, Writing – Review & Editing

<u>Anna G. Anders</u>: Investigation, Writing – Review & Editing

<u>Brandon T. Ruotolo</u>: Writing – Review & Editing <u>Robert T. Kennedy</u>: Writing – Review & Editing

<u>Kristin S. Koutmou</u>: Conceptualization, Writing – Review & Editing

## ACKNOWLEDGMENTS

We would like to acknowledge Dr. B. Shay and Dr. M. Sorenson for their thoughtful discussions regarding LC-MS/MS method development and sample preparation. We thank the following funding sources for their support: National Institutes of Health (NIGMS R35 GM128836), National Science Foundation (NSF CAREER 2045562 to K.S.K., GFRP to J.D.J., and NSF CHE 1904146 to R.T.K.),and Research Corporation for Science Advancement (Cottrell Scholar Award to K.S.K.)

## ABBREVIATIONS

(m^1^G): 1-methylguanosine
(m^2^G): N2-methylguanosine
(m^2^_2_G): N2, N2-dimethylguanosine
(m^5^U): 5-methyluridine
(Ψ): pseudouridine
(m^1^Ψ): N1-methylpseudouridine
(Am): 2’O-methyladenosine
(Um): 2’O-methyluridine
(Cm): 2’O-methylcytidine
(Gm): 2’O-methylguanosine
(m^7^G): N7-methylguanosine
(m^1^A): 1-methyladenosine
(DHU): dihy-drouridine
(ac^4^C): N4-actetylcytidien
(m^3^C): 3-methylcyti-dine
(i^6^A): isopentenyl-N6-adenosine
(m^6^A): N6-methyl-adenosine
(8-oxo-G): 8-oxoguanosine

## SYNOPSIS TOC

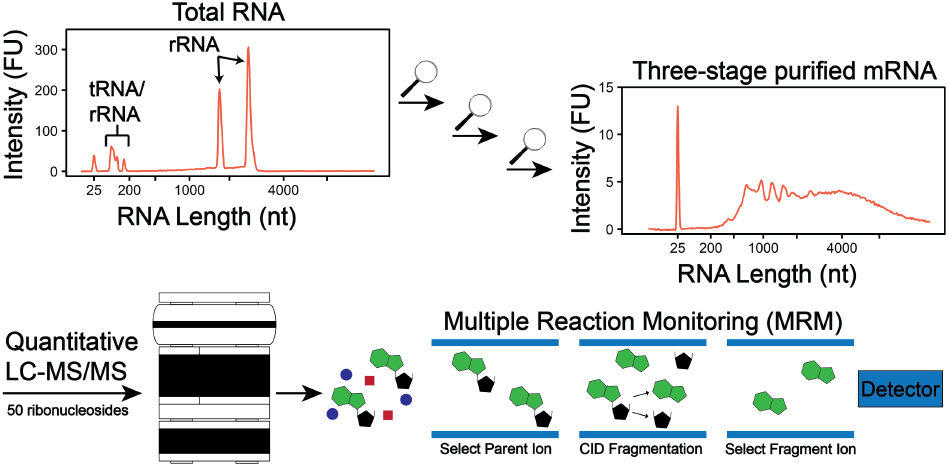

