## Supplementary material for "Methylated guanosine and uridine modifications in *S. cerevisiae* mRNAs modulate translation elongation": Jones_etal_Supplemental Figures

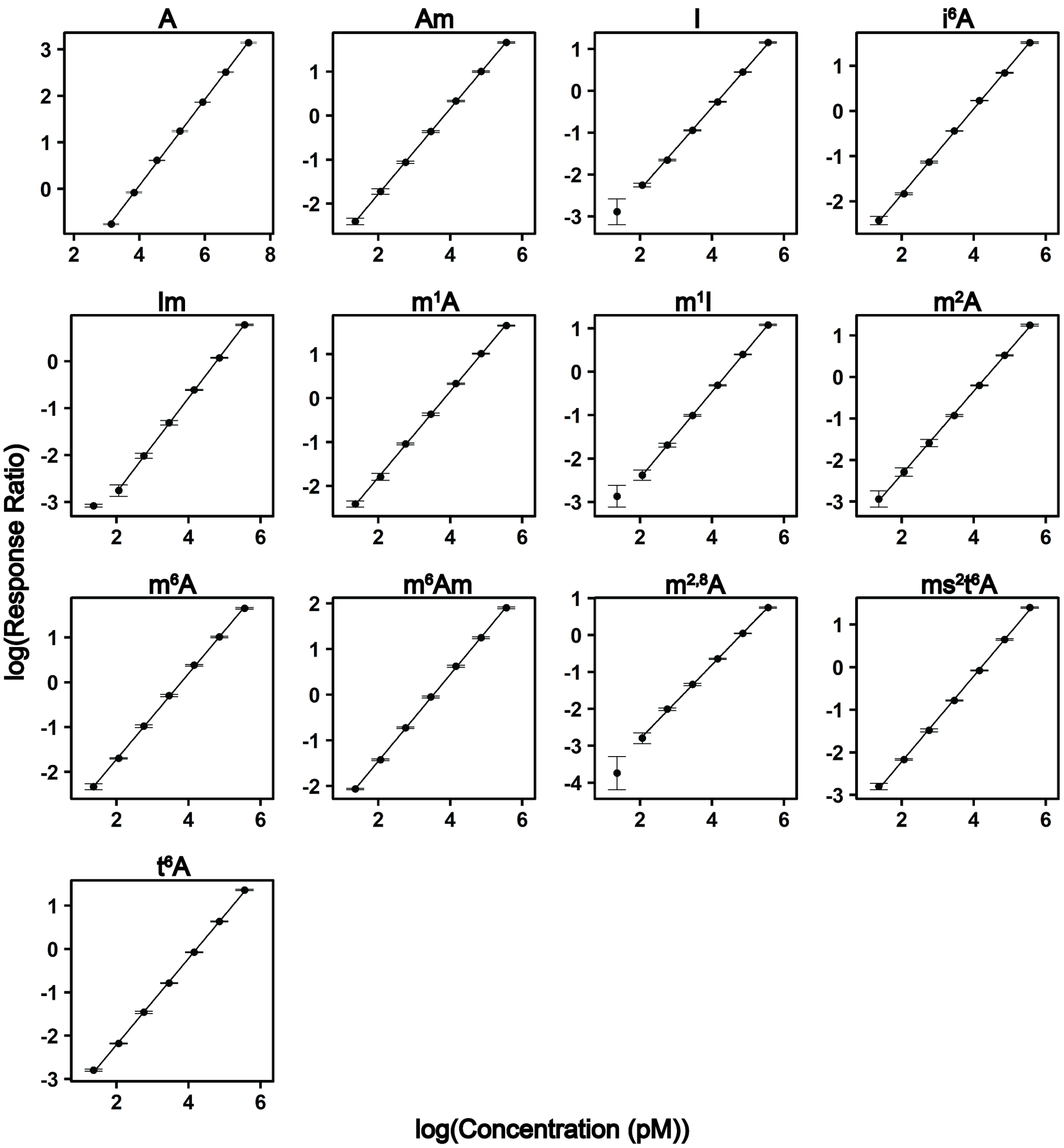


**Supplemental Figure S1: Calibration curves used to quantify adenosine modification concentrations.** Calibration curves of adenosine ribonucleoside modifications plotted in log(response ratio) vs. log(concentration (pM)). The linear regression, limit of detection, and $R^2$ are displayed in **Supplemental Table S1.**

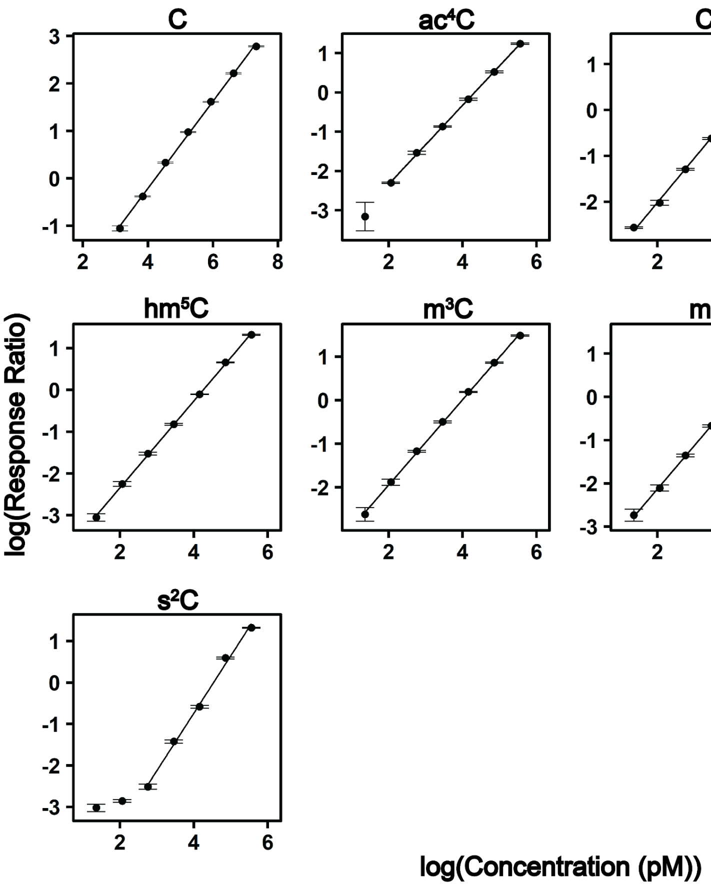


**Supplemental Figure S2: Calibration curves used to quantify cytidine modification concentrations.** Calibration curves of cytidine ribonucleoside modifications plotted in log(response ratio) vs. log(concentration (pM)). The linear regression, limit of detection, and $R^2$ are displayed in **Supplemental Table S1.**

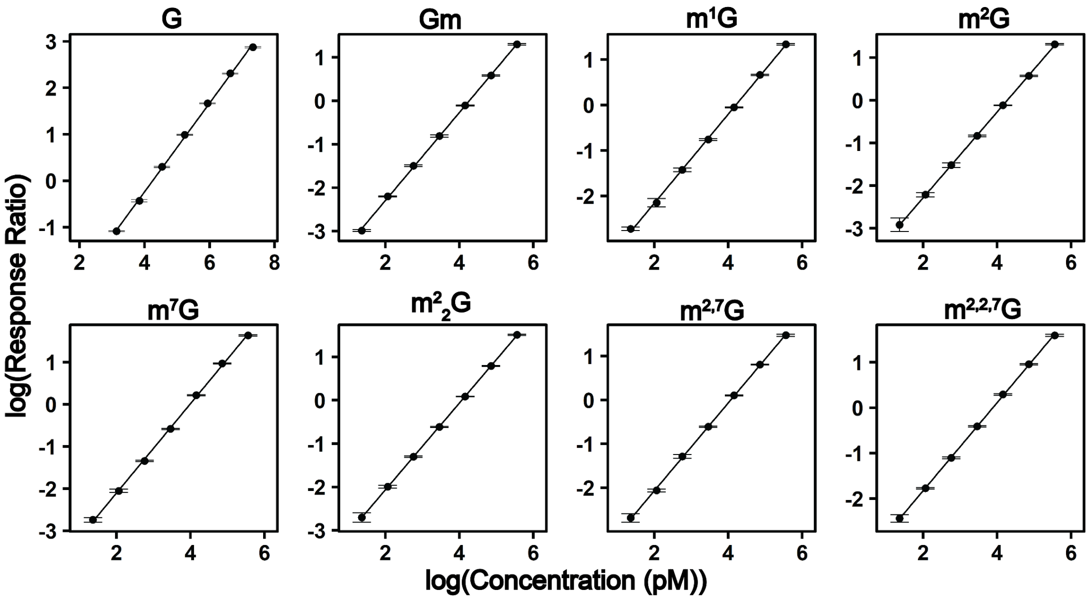


**Supplemental Figure S3: Calibration curves used to quantify guanosine modification concentrations.** Calibration curves of guanosine ribonucleoside modifications plotted in log(response ratio) vs. log(concentration (pM)). The linear regression, limit of detection, and $R^2$ are displayed in **Supplemental Table S1.**

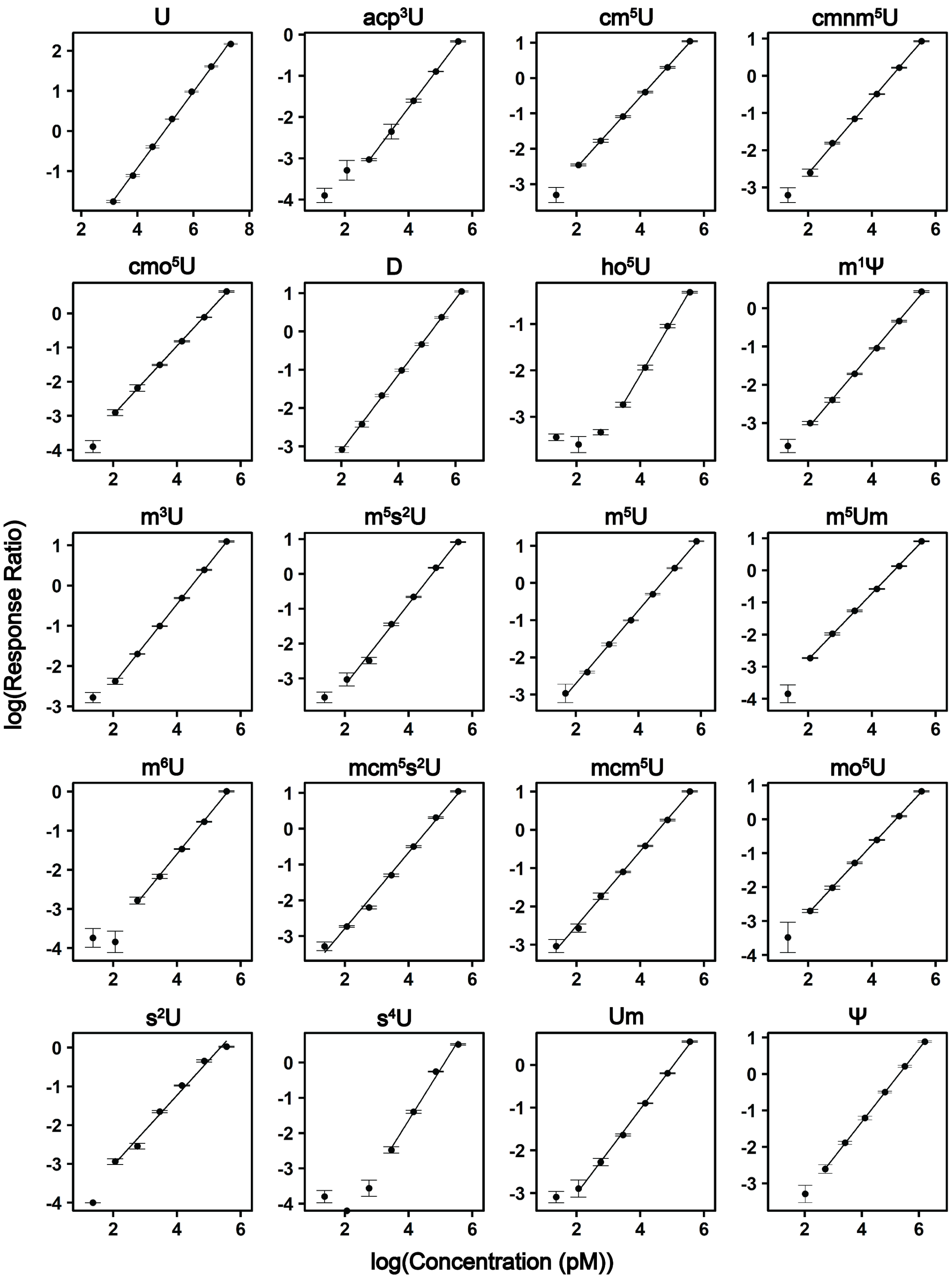


**Supplemental Figure S4: Calibration curves used to quantify uridine modification concentrations.** Calibration curves of uridine ribonucleoside modifications plotted in log(response ratio) vs. log(concentration (pM)). The linear regression, limit of detection, and $R^2$ are displayed in **Supplemental Table S1.**

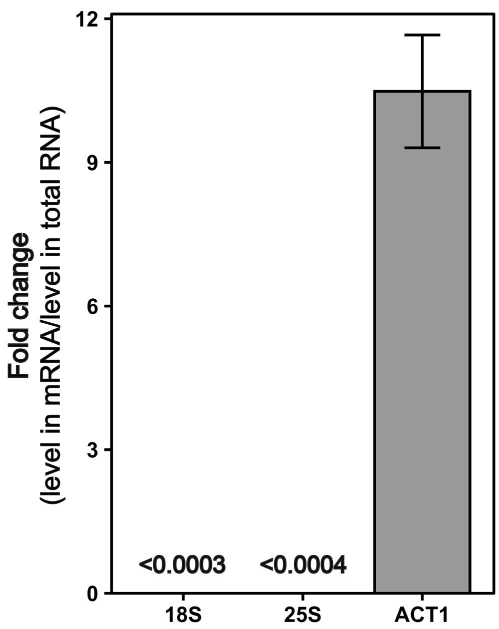


**Supplemental Figure S5: Ribosomal RNAs are depleted in three-stage purified mRNA.** qRT-PCR demonstrates that the 18S and 25S rRNAs are depleted by greater than 3000-fold in the purified mRNA. Contrarily, ACT1 is enriched by greater than 10-fold. This data in addition to the Bioanalyzer electropherograms, RNA-seq, and LC-MS/MS proves that our three-stage purified mRNA is highly pure.

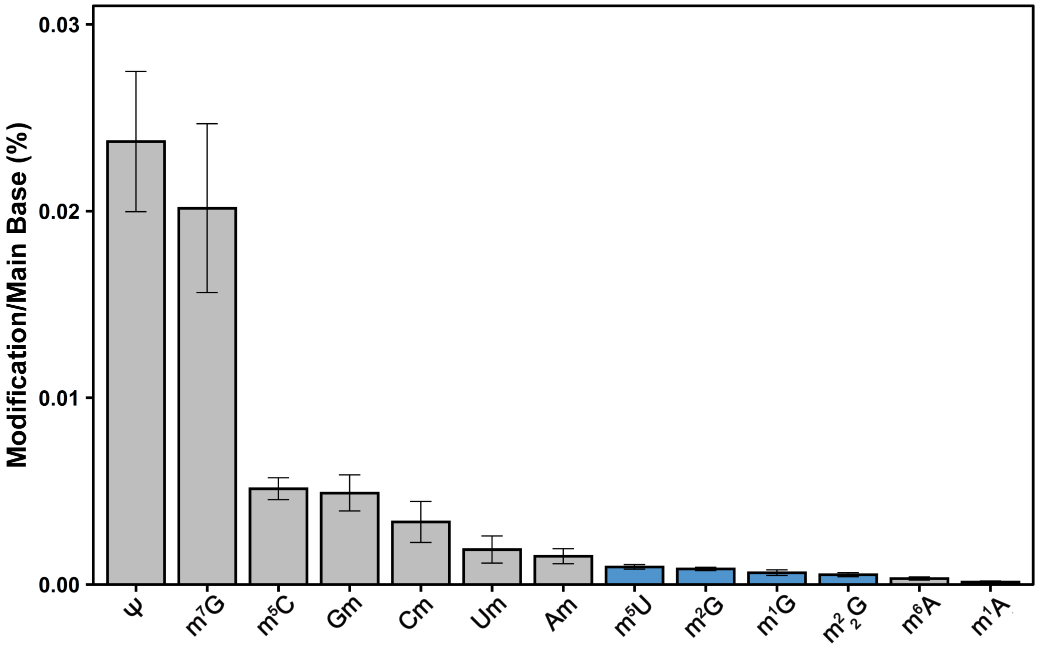


**Supplemental Figure S6: Ribonucleoside modification abundance in the three-stage purified mRNA.** The ribonucleoside abundance is represented as modification/main base% (i.e., $m^7G/G\%$) where pseudouridine was the most abundant modification detected. All modifications detected were previously detected in purified mRNA besides for the three methylated guanosine modifications displayed in blue ($m^1G$, $m^2G$, and $m^2{}_2G$). Our improvements regarding LC-MS/MS sensitivity and mRNA purity enables us to confidently claim these modifications exist with *S. cerevisiae* mRNA.

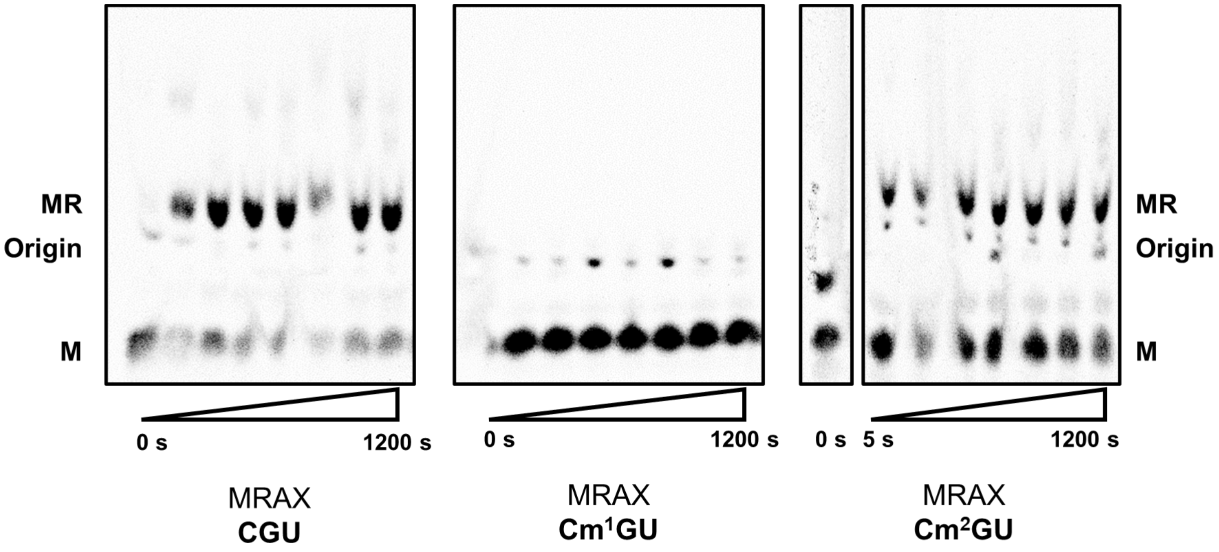


**Supplemental Figure S7:** Electrophoretic TLC displaying the translation products of CGU, $Cm^1GU$, and $Cm^2GU$ codons in the presence of arginine tRNA (ArgTC), forming MR dipeptide over the span of 1200 seconds.

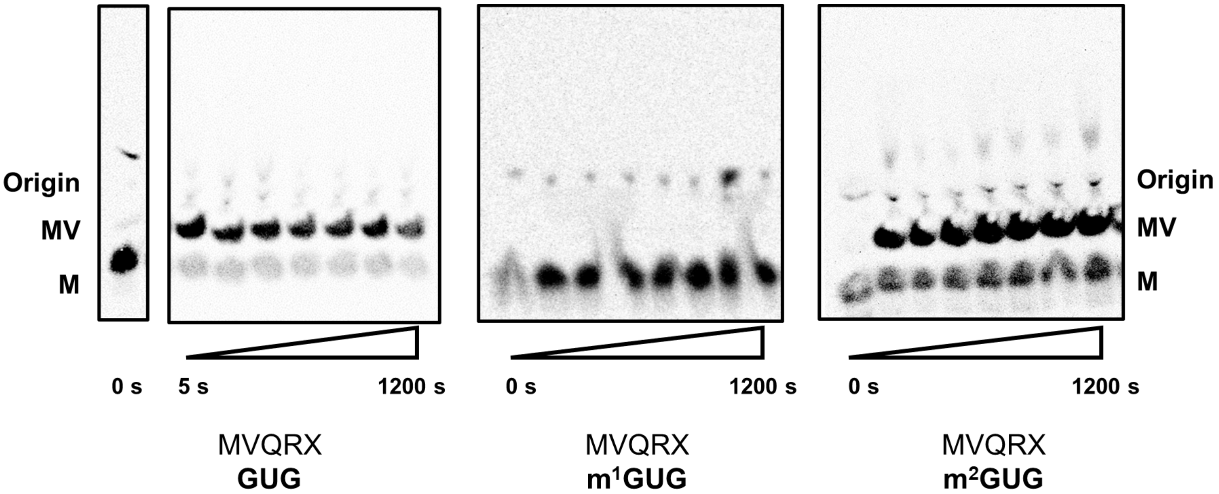


**Supplemental Figure S8:** Electrophoretic TLC displaying the translation products of GUG, $m^1$GUG, and $m^2$GUG codons in the presence of valine tRNA (ValTC), forming MV dipeptide over the span of 1200 seconds.

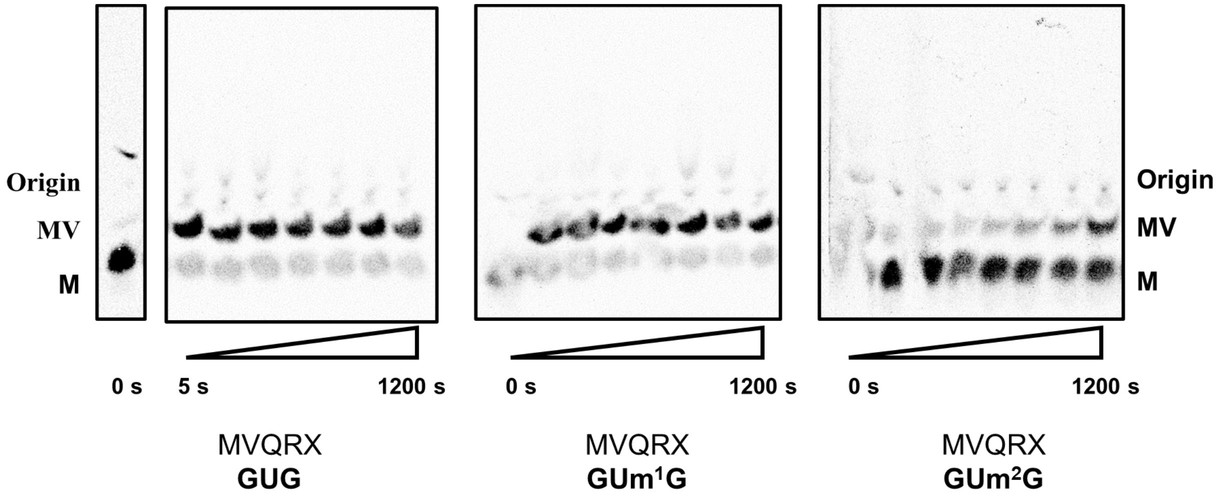


**Supplemental Figure S9:** Electrophoretic TLC displaying the translation products of GUG, GUm$^1$G, and GUm$^2$G codons in the presence of valine tRNA (ValTC), forming MV dipeptide over the span of 1200 seconds.

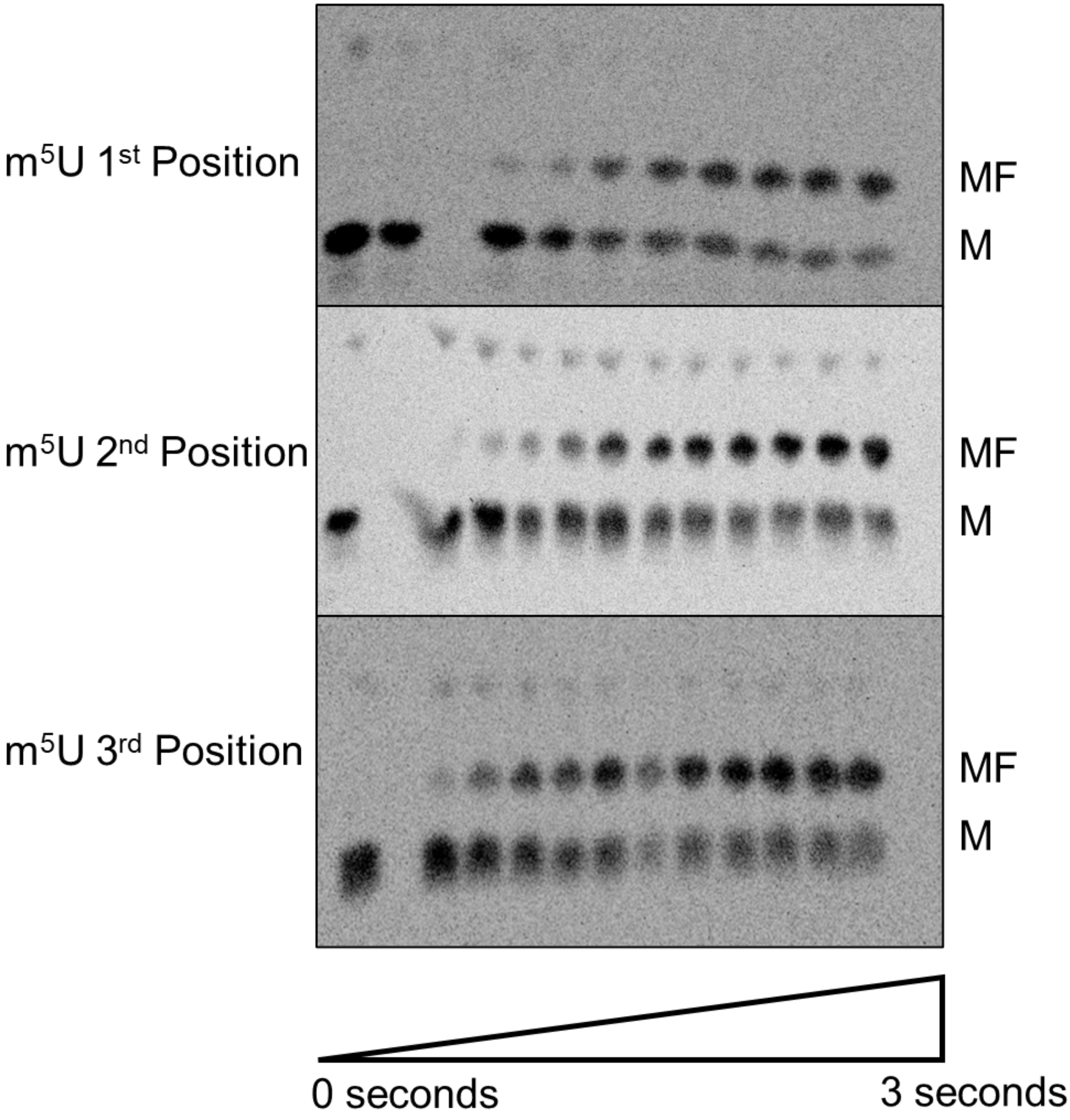


**Supplemental Figure S10:** Electrophoretic TLC displaying the translation products of $m^5U$ messages in the presence of phenylalanine tRNA (PheTC), forming MF dipeptide over the span of 3 seconds.

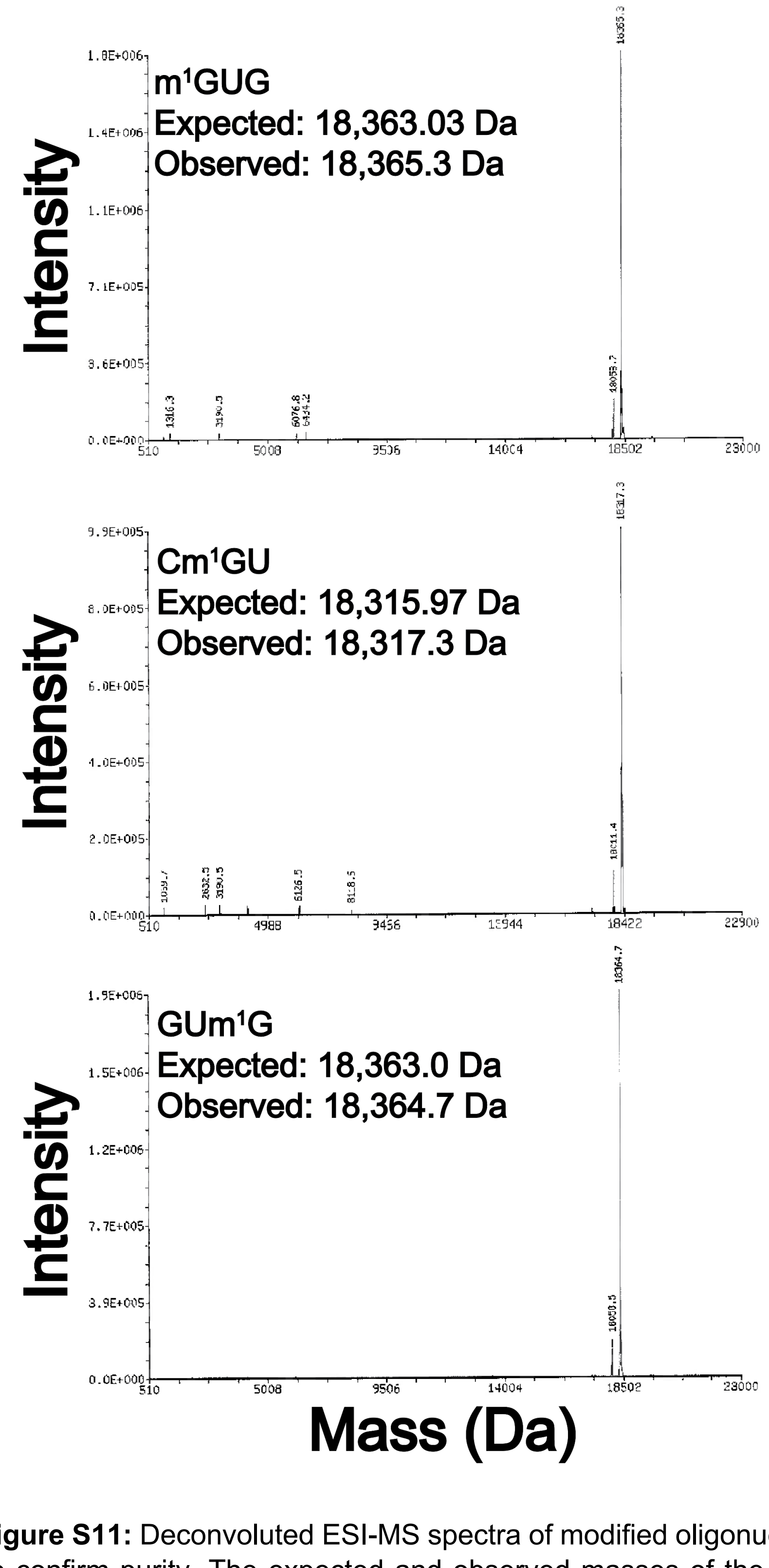


**Supplemental Figure S11:** Deconvoluted ESI-MS spectra of modified oligonucleotides provided by Dharmacon to confirm purity. The expected and observed masses of the m$^1$GUG, Cm$^1$GU, and GUm$^1$G modified codon oligonucleotides are found in the top, middle, and bottom panels, respectively. Minor n-1 oligonucleotides products were detected, but they would not affect the *in vitro* translation assays because the nucleotide loss occurs in the non-coded region of the purchased mRNA transcript.

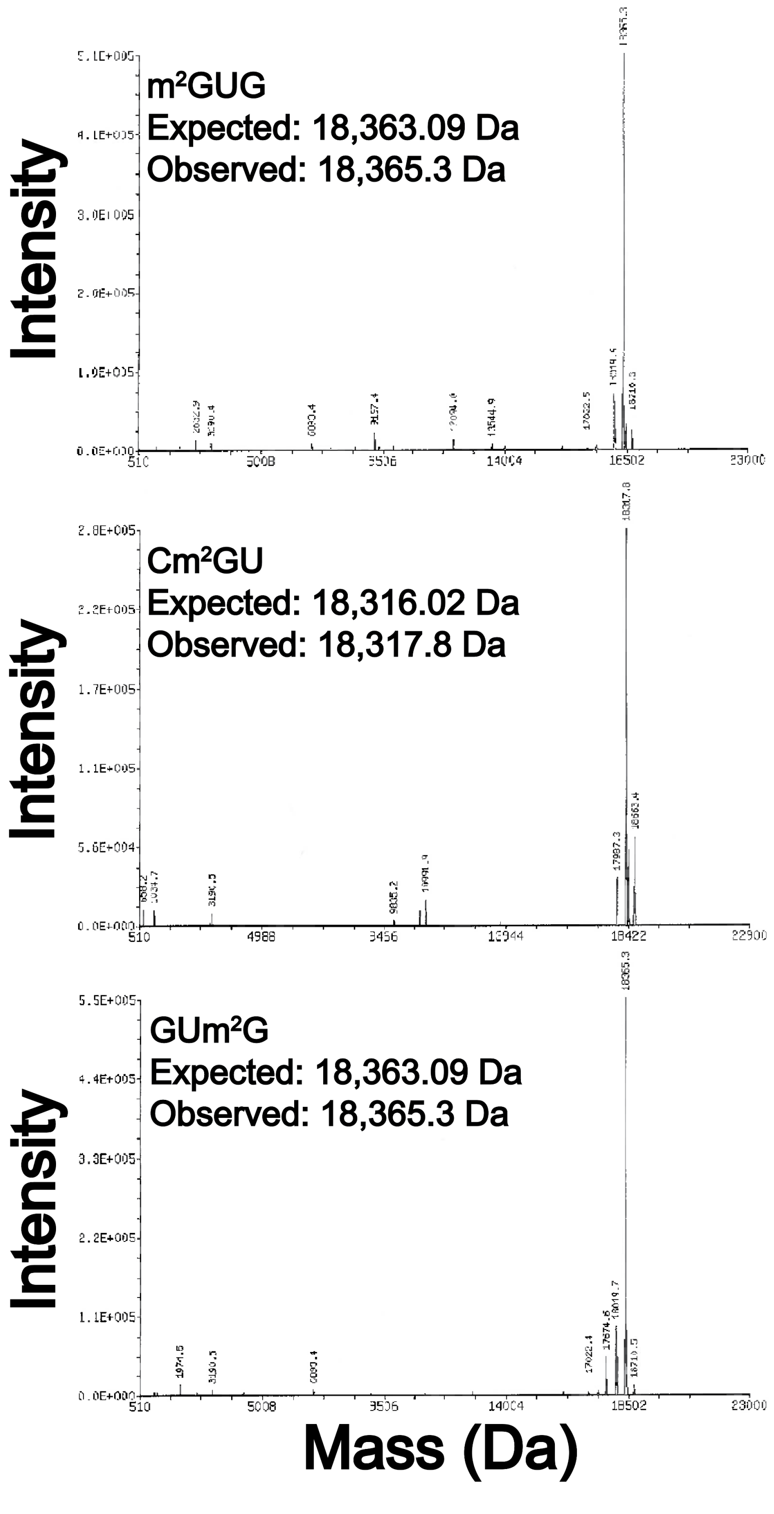


**Supplemental Figure S12:** Deconvoluted ESI-MS spectra of modified oligonucleotides provided by Dharmacon to confirm purity. The expected and observed masses of the m$^2$GUG, Cm$^2$GU, and GUm$^2$G modified codon oligonucleotides are found in the top, middle, and bottom panels, respectively. Minor n-1 oligonucleotides products were detected, but they would not affect the *in vitro* translation assays because the nucleotide loss occurs in the non-coded region of the purchased mRNA transcript.

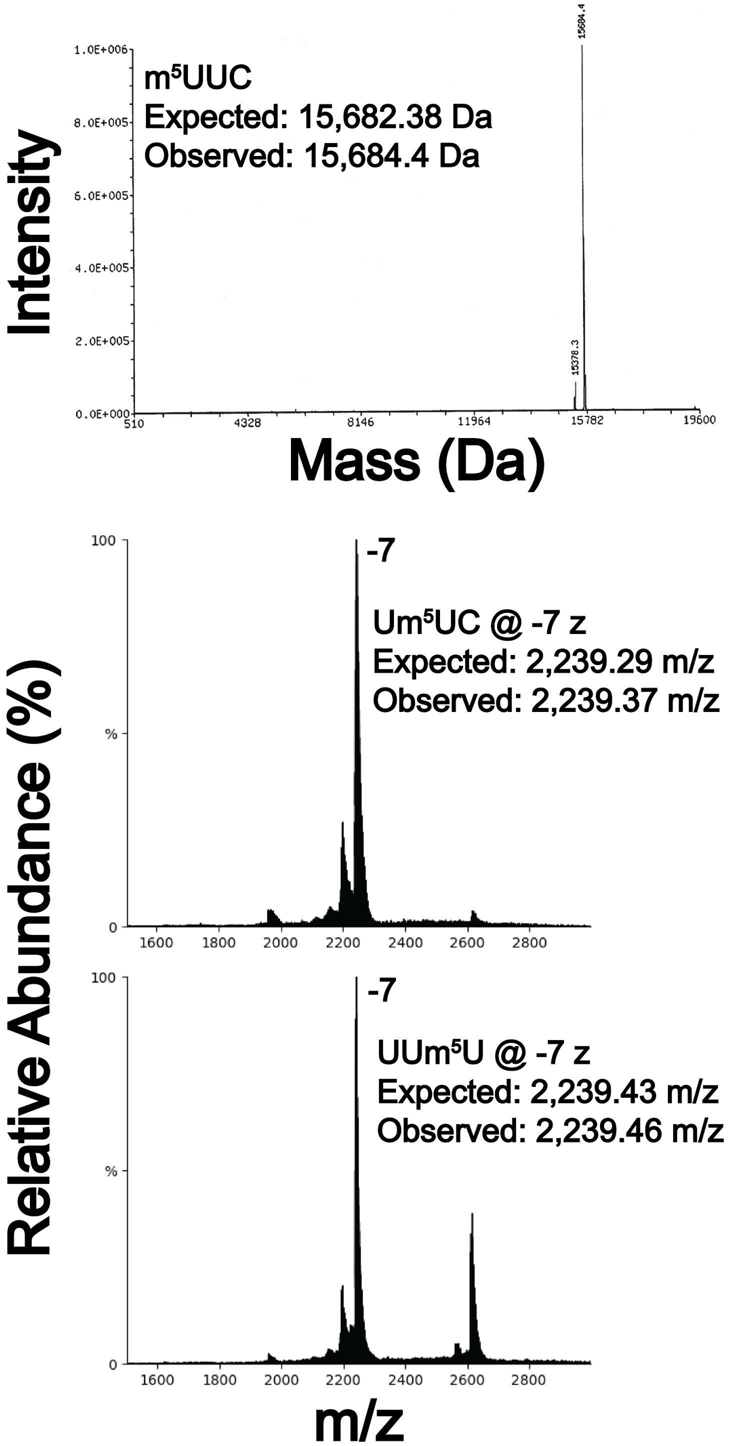


**Supplemental Figure S13:** Deconvoluted ESI-MS spectra of m$^5$UUC modified codon oligonucleotides provided by Dharmacon to confirm purity (top panel). Full scan spectra of Um$^5$UC (middle) and UUm$^5$U (bottom) modified codon oligonucleotide. The corresponding expected and observed mass (Da) or mass-to-charge (m/z) is displayed for each spectrum. Minor n-1 oligonucleotides products were detected, but they would not affect the *in vitro* translation assays because the nucleotide loss occurs in the non-coded region of the purchased mRNA transcript.
